# Vitamin B2 metabolism promotes FSP1 stability to prevent ferroptosis

**DOI:** 10.1101/2025.08.05.668752

**Authors:** Kirandeep K. Deol, Cynthia A. Harris, Sydney J. Tomlinson, Cody E. Doubravsky, Alyssa J. Mathiowetz, James A. Olzmann

**Author notes:** Address correspondence to: James A. Olzmann.

## Abstract

Ferroptosis, a regulated form of cell death driven by excessive lipid peroxidation, has emerged as a promising therapeutic target in cancer. Ferroptosis suppressor protein 1 (FSP1) is a critical regulator of ferroptosis resistance, yet the mechanisms controlling its expression and stability remain mostly unexplored. To uncover regulators of FSP1 abundance, we conducted CRISPR-Cas9 screens utilizing a genome-edited, dual-fluorescent FSP1 reporter cell line, identifying both transcriptional and post-translational mechanisms that determine FSP1 levels. Notably, we identified riboflavin kinase (RFK) and FAD synthase (FLAD1), enzymes which are essential for synthesizing flavin adenine dinucleotide (FAD) from vitamin B2, as key contributors to FSP1 stability. Biochemical and cellular analyses revealed that FAD binding is critical for FSP1 activity. FAD deficiency, and mutations blocking FSP1-FAD binding, triggered FSP1 degradation via a ubiquitin-proteasome pathway that involves the E3 ligase RNF8. Unlike other vitamins that inhibit ferroptosis by scavenging radicals, vitamin B2 supports ferroptosis resistance through FAD cofactor binding, ensuring proper FSP1 stability and function. This study provides a rich resource detailing mechanisms that regulate FSP1 abundance and highlights a novel connection between vitamin B2 metabolism and ferroptosis resistance with implications for therapeutic strategies targeting FSP1 in cancer.

## INTRODUCTION

Ferroptosis is a regulated form of cell death driven by the iron-dependent accumulation of oxidatively-damaged polyunsaturated fatty acid (PUFA)-containing membrane phospholipids (i.e., lipid peroxides)^1–5^. The primary defense against ferroptosis is mediated by glutathione peroxidase 4 (GPX4), which reduces lipid hydroperoxides to non-toxic alcohols using glutathione as a cofactor^6^. In some cases, inhibition of glutathione synthesis or GPX4 triggers uncontrolled lipid peroxidation, culminating in ferroptotic cell death involving plasma membrane rupture^2,3^. Given its unique mechanism, ferroptosis has emerged as a promising target for therapeutic intervention, particularly in cancer, where resistance to other forms of cell death often underlies treatment failure^7,8^. Indeed, many cancer cells exhibit an inherent susceptibility to ferroptosis due to their altered metabolic states, iron levels, and membrane composition, such as mesenchymal cancer cells that give rise to therapy-resistant metastatic tumors^9–12^. Exploiting these vulnerabilities with ferroptosis-inducing agents has demonstrated promise in preclinical models^7,8^, though it is still unclear whether ferroptosis-targeting therapeutics will be effective in the clinic.

A second cellular defense against ferroptosis involves the generation and recycling of radical trapping antioxidants (RTAs)^1,2^. A key player in this mechanism is ferroptosis suppressor protein 1 (FSP1), which is a myristoylated, membrane-bound oxidoreductase for non-mitochondrial ubiquinone (coenzyme Q10, CoQ10)^13,14^ and vitamin K^15,16^, both of which act as RTAs to prevent lipid peroxidation. FSP1’s activity complements GPX4, providing a parallel protective pathway to suppress lipid peroxidation and ferroptosis. This redundancy underscores FSP1’s importance in ferroptosis resistance, particularly under conditions in which GPX4 is inhibited. Knockout (KO) of *FSP1*^7,8,17^ and small molecule inhibition of FSP1^14,18–20^ sensitize diverse cancer cell lines to ferroptosis inducers. *FSP1* KO also suppresses tumor growth in xenograft models^13,17,20,21^. Moreover, FSP1 expression strongly correlates with ferroptosis resistance^13,14^ and high FSP1 expression is linked to poor outcomes in cancer^17^, further highlighting its potential as a therapeutic target. Despite its relevance, little is known about the mechanisms that govern FSP1 expression and stability, leaving a critical gap in our understanding of how FSP1 is regulated.

In this study, we address this knowledge gap by employing a series of CRISPR-Cas9 genetic screens in a genome-edited, dual-fluorescent FSP1 reporter cell line. Our findings reveal and stratify transcriptional and post-translational regulators of FSP1 abundance, shedding light on the mechanisms that control its expression and stability. Among these, we identify riboflavin kinase (RFK) and FAD synthase (FLAD1), enzymes essential for the biosynthesis of flavin adenine dinucleotide (FAD) from vitamin B2, as critical contributors to FSP1 stability and function. These findings uncover a link between vitamin B2 metabolism and ferroptosis resistance, emphasizing the unique role of FAD in supporting FSP1-mediated lipid antioxidant defenses. This work also provides a comprehensive resource on FSP1 regulatory mechanisms and introduces new opportunities for targeting ferroptosis in cancer therapy.

## RESULTS

### Generation of a genome-edited FSP1 GFP-P2A-BFP reporter cell line

CRISPR-Cas9 genome editing was employed to generate a fluorescence-based FSP1 reporter cell line in U-2 OS osteosarcoma cells. U-2 OS cells express high levels of FSP1 and rely on FSP1 to prevent ferroptosis^13^. Endogenous tagging enables analysis of expression and avoids artifacts associated with ectopic overexpression. In the resulting FSP1^GFP-P2A-BFP^ cell line, FSP1 was endogenously tagged at its carboxyl terminus with green fluorescent protein (GFP), followed by a P2A sequence and blue fluorescent protein (BFP) (**Fig. 1A**). The P2A sequence encodes a short peptide that induces ribosomal skipping, enabling translation of BFP from the same transcript as FSP1-GFP (**Fig. 1A**). This design facilitates multiple analyses: GFP levels report on FSP1 protein abundance, BFP levels reflect *FSP1* transcription, and the GFP:BFP ratio provides insights into the post-translational stability of FSP1.

**Fig. 1.**
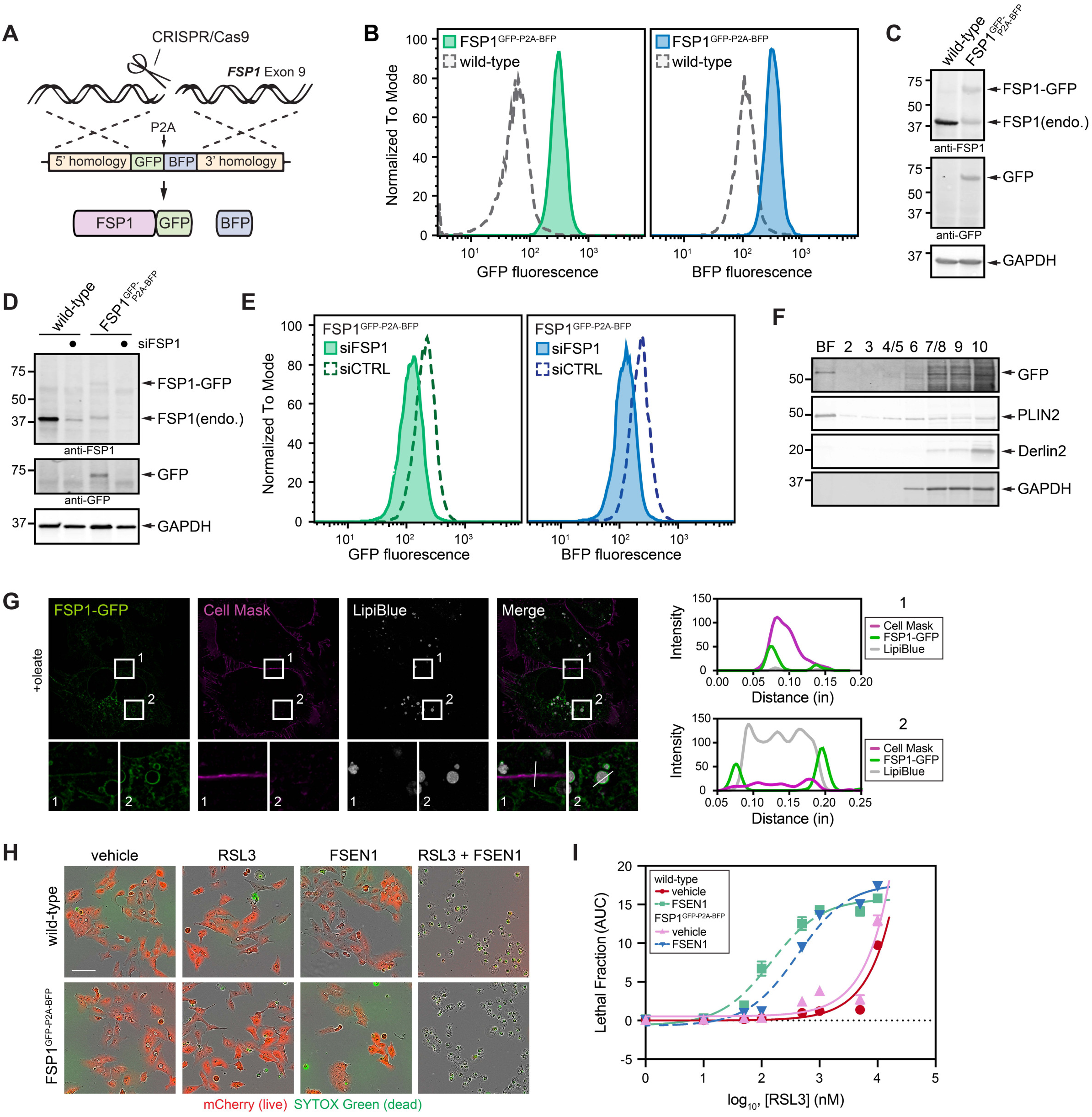
Construction of a genome-edited fluorescent reporter for monitoring FSP1 levels in human cells. **a**, Schematic of CRISPR-Cas9 strategy used to generate the FSP1^GFP-P2A-BFP^ reporter system. **b**, Fluorescence histograms of wild-type control and genomic FSP1^GFP-P2A-BFP^ knock-in cells. **c**, Immunoblot of lysates from FSP1^GFP-P2A-BFP^ cells. endo., endogenous FSP1. **d**, Immunoblot of control and FSP1^GFP-P2A-BFP^ cells incubated with siRNAs against FSP1 for 72 h. **e**, Fluorescence histograms of FSP1^GFP-P2A-BFP^ cells from (**d**) including siRNAs from scramble control. **f**, Buoyant fractions, enriched with lipid droplets from FSP1^GFP-P2A-BFP^ cells, were isolated by sucrose gradient fractionation and analyzed by western blot. *BF, buoyant fraction*. **g**, FSP1^GFP-P2A-BFP^ subcellular distribution by live-cell microscopy. Cells were incubated with 200 µM oleate to induce lipid droplet formation, 500 nM LipiBlue to label lipid droplets, and 5 µg/mL CellMask Deep Red to label the plasma membrane. Line intensity plots showing colocalization between FSP1^GFP-P2A-BFP^ and organelle markers (right). **h**, Live-cell imaging of mCherry-expressing control and FSP1^GFP-P2A-BFP^ cells incubated with SYTOX Green and treated with 100 nM RSL3 for 24 h. Scale bar, 100 µm. **i**, Dose response of RSL3-induced cell death of wild-type and FSP1^GFP-P2A-BFP^ reporter cells in the presence of 2 µM FSEN1.

The FSP1^GFP-P2A-BFP^ cell line was readily distinguishable from wild-type control cells by flow cytometry, displaying a single peak of GFP and BFP fluorescence (**Fig. 1B, Extended Data Fig. S1A**). Western blot analysis indicated the presence of both endogenous, untagged FSP1 and GFP-tagged FSP1 in FSP1^GFP-P2A-BFP^ cells (**Fig. 1C**). Anti-GFP antibodies confirmed a band at the expected molecular weight for FSP1-GFP (∼67.5 kDa; **Fig. 1C**). Correct in-frame insertion of GFP was also confirmed by DNA sequencing (**Extended Data Fig. S1B**). Knockdown of FSP1 using FSP1-targeting siRNAs resulted in decreased levels of both endogenous FSP1 and FSP1-GFP by western blotting (**Fig. 1D**) and in a corresponding reduction in GFP and BFP fluorescence by flow cytometry (**Fig. 1E**). These results indicate that the FSP1^GFP-P2A-BFP^ reporter cell line is heterozygous, expressing both untagged FSP1 and FSP1-GFP, and that FSP1-GFP and BFP are translated from the same transcript.

Consistent with previous observations of FSP1 localization to lipid droplets^13,14^, sucrose gradient biochemical fractionation revealed that a portion of FSP1-GFP associates with buoyant, lipid droplet-enriched fractions containing PLIN2 (**Fig. 1F**). Additionally, FSP1 is known to localize to other membranes, including the plasma membrane^13^. Super-resolution fluorescence imaging showed FSP1-GFP partially co-localizes with the plasma membrane marker CellMask and encircles LipiBlue-stained lipid droplets (**Fig. 1G**). To assess the functionality of FSP1 in the reporter cell line, we measured ferroptosis sensitivity in wild-type and FSP1^GFP-P2A-BFP^ cells. FSP1^GFP-P2A-BFP^ cells exhibited similar resistance to RSL3 as wild-type cells, with both showing strong sensitization to RSL3-induced ferroptosis following FSEN1 treatment (**Fig. 1H, I, Extended Data Fig. S1C**). Finally, we examined the impact of proteolysis inhibitors to determine if FSP1 is actively degraded. These included inhibitors of autophagy (3-methyladenine), lysosomal acidification (BafA1), p97/VCP (CB5083), ubiquitination (MLN7243), and proteasomal activity (MG132). BFP levels remained unaffected by these treatments (**Extended Data Fig. S1D**). However, GFP levels and the GFP:BFP fluorescence ratio showed a small but significant increase in response to CB5083, MLN7243, and MG132, indicating that FSP1 is relatively stable but a portion is actively degraded by the ubiquitin-proteasome system (**Extended Data Fig. S1D-F**).

Collectively, these results demonstrate that FSP1-GFP is correctly localized and functional, and that the FSP1^GFP-P2A-BFP^ cell line provides a robust, fluorescence-based reporter system for investigating FSP1 regulation.

### Genome-wide genetic screen identifies regulators of FSP1 levels

To globally identify the genetic factors that govern FSP1 levels, we conducted a genome-wide CRISPR-Cas9 loss-of-function screen where Cas9-expressing FSP1^GFP-P2A-BFP^ cells were transduced with a sgRNA library (∼10 sgRNAs per gene) and sorted into GFP^high^ and GFP^low^ populations (**Fig. 2A**). Following FACS, the sgRNA frequencies in GFP^high^ and GFP^low^ cells were compared to generate an adjusted log_2_ enrichment value (gene effect), a confidence metric calculated from the log-likelihood ratio (gene score), and a *p*-value for each gene using Cas9 high-Throughput maximum Likelihood Estimator (casTLE) ^22,23^. At a 10% false discovery rate (FDR), we identified 635 regulators of FSP1, including 498 genes whose depletion increased FSP1 levels and 137 genes whose depletion reduced FSP1 levels (**Fig. 2B, Table S1**). As expected, *FSP1* itself emerged as a high confidence hit, with 9 out of 10 sgRNAs showing depletion from the isolated GFP^high^ cells and enrichment in the GFP^low^ cells (**Fig. 2C**). Unbiased gene ontology analyses revealed significant enrichment in several functional categories, including selenocysteine incorporation, redox homeostasis, ubiquinone biosynthesis, the ubiquitin-proteasome system, and other cellular processes (**Fig. 2D**).

**Fig. 2.**
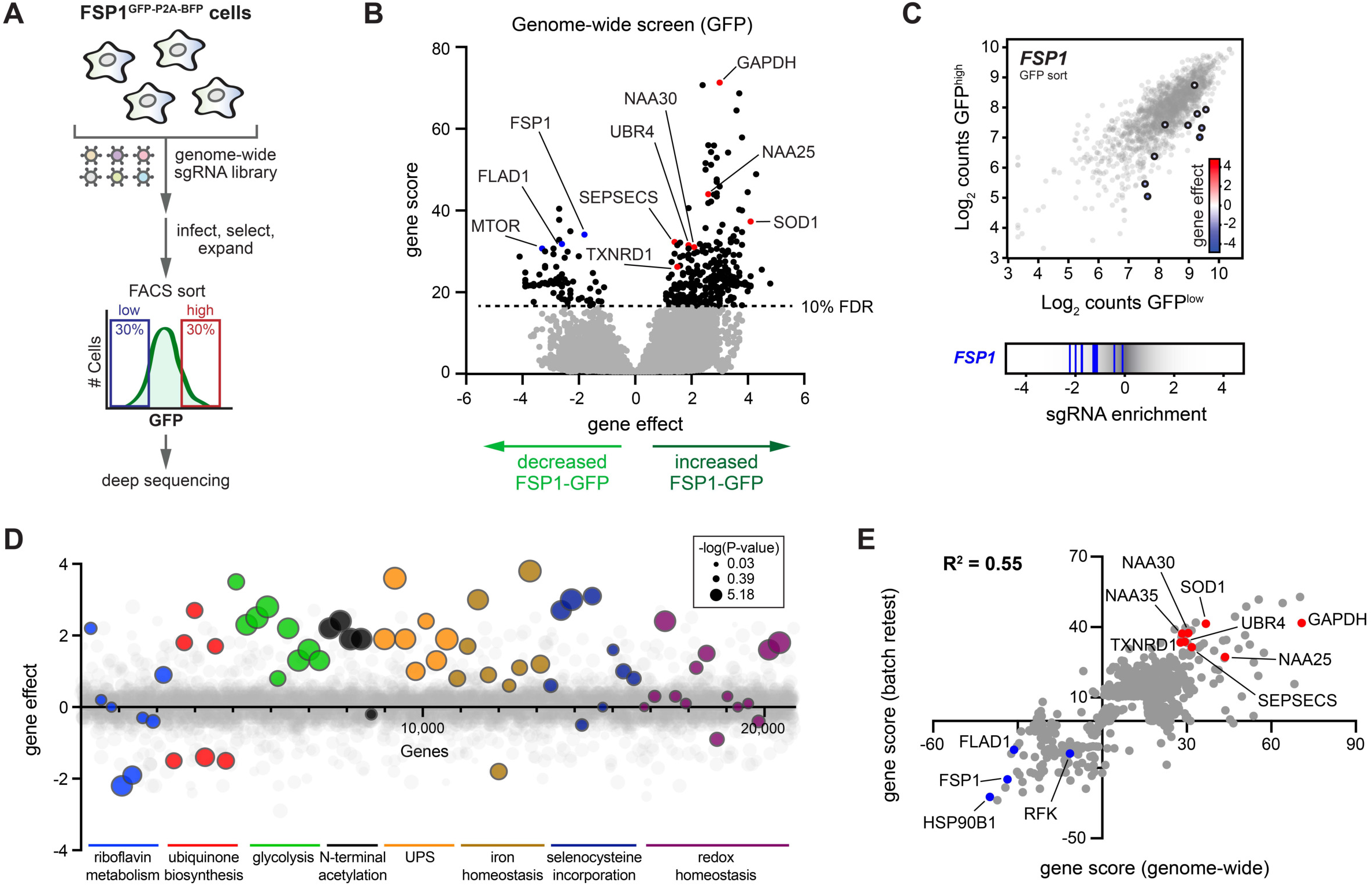
Genome-wide CRISPR analysis reveals regulators of FSP1 abundance. **a**, Schematic of the genome-wide CRISPR-Cas9 screening strategy. **b**, Gene effects and gene scores calculated for individual genes analyzed in the genome-wide CRISPR screen. **c**, Cloud plot indicating count numbers corresponding to *FSP1* (color scale) and control (gray scale) sgRNAs (top). Frequency histogram indicating the distribution of the relative enrichment for *FSP1* sgRNAs (blue lines, bottom). The gray line indicates the mean of the control sgRNA distribution. **d**, Bubble plot of the genes identified from the genome-wide screen with a 10% FDR cutoff, where each bubble color represents an enriched GO term functional cluster. **e**, Scatter plot of signed gene scores for individual genes from genome-wide screen (x-axis) versus batch retest screen (y-axis).

To rigorously validate our screen results and reduce false positives^24–27^, we conducted duplicate secondary pooled screens (also known as batch retest screens) at high coverage using a custom-synthesized sgRNA library. This cloned library included the 635 regulatory genes identified at a 10% FDR from the genome-wide screen as well as 100 manually curated ferroptotic genes of interest and maintained excellent diversity (**Extended Data Fig. S2A**). FSP1^GFP-P2A-BFP^ cells expressing Cas9 were transduced with the custom batch retest sgRNA library and FACS was used to sort GFP^high^ and GFP^low^ populations, replicating the approach from the initial screen. The results closely aligned with those of the genome-wide screen (R² = 0.55), reaffirming the identification of key regulatory candidates across diverse functional categories (**Fig. 2E, Table S2**). Together, the genome-wide and batch retest screens provide a robust and comprehensive view of the regulatory landscape governing FSP1 levels in U-2 OS osteosarcoma cells.

### Parallel genetic screens reveal transcriptional and post-translational regulators of FSP1

Our genome-wide and batch retest screens that used GFP as a reporter identified regulators of FSP1 abundance that likely act through both transcriptional and post-translational regulation. To systematically differentiate these mechanisms, we performed additional genetic screens leveraging BFP as a reporter for *FSP1* transcription and the GFP:BFP ratio as a reporter for FSP1 post-translational stability (**Fig. 3A**). Duplicate high-coverage screens were conducted in FSP1^GFP-P2A-BFP^ cells expressing Cas9 using our custom batch retest sgRNA library. To identify transcriptional regulators of FSP1, cells with BFP^high^ and BFP^low^ fluorescence were isolated by FACS. To identify post-translational regulators of FSP1 stability, cells with GFP:BFP^high^ and GFP:BFP^low^ ratio were similarly isolated by FACS. Recognizing that lipid peroxidation and ferroptosis might influence FSP1 expression and stability, we performed these screens under both basal conditions and RSL3 treatment (100 nM), a dose that induces lipid peroxidation but minimal cell death (**Extended Data Fig. S2B**).

**Fig. 3.**
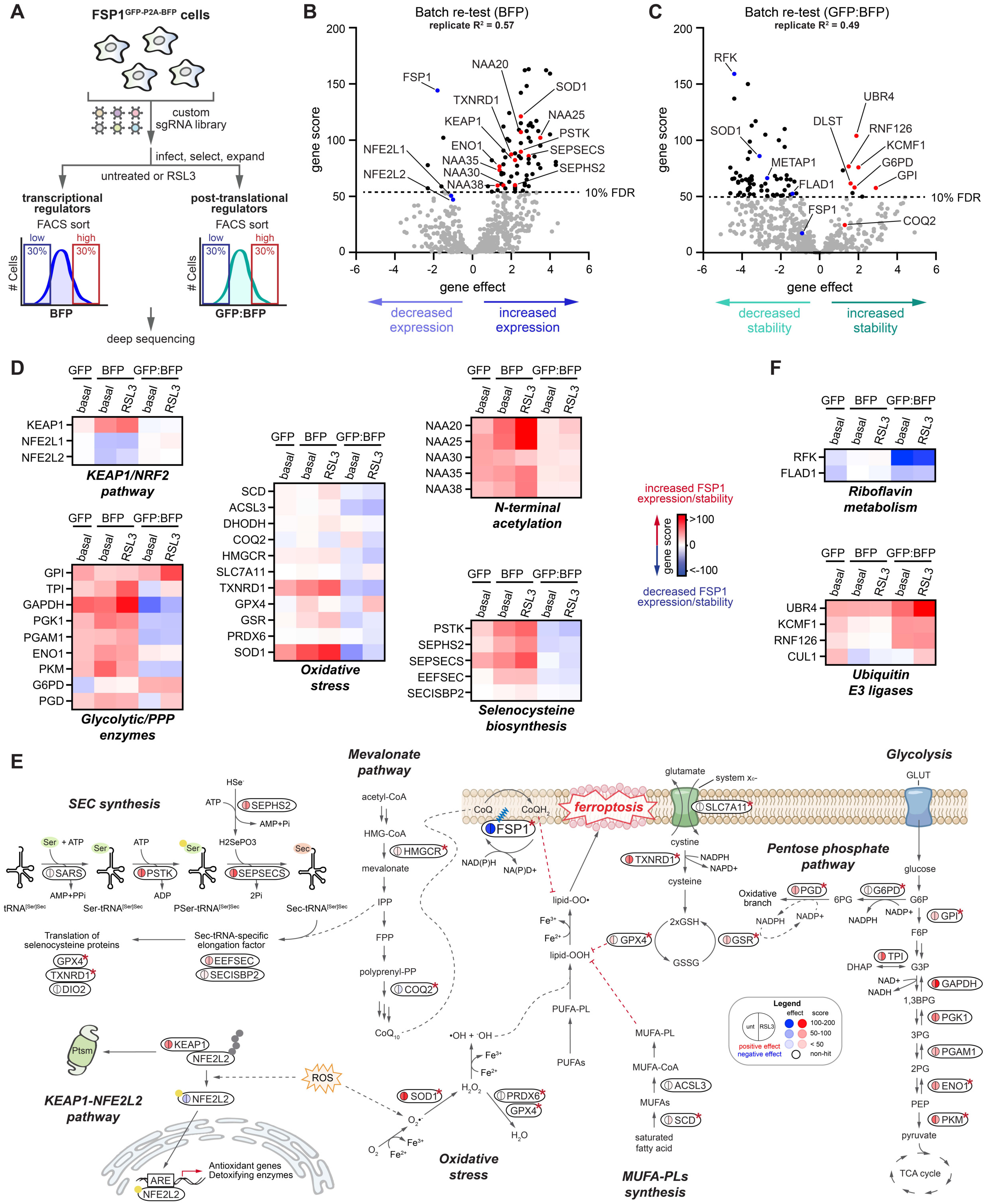
Parallel CRISPR-Cas9 screens uncouple regulators of FSP1 expression and stability. **a**, Schematic of the CRISPR-Cas9 batch retest screening strategy. **b-c**, Gene effects and gene scores calculated for individual genes analyzed in the batch retest CRISPR screens for expression (BFP, **b**) or stability (GFP:BFP, **c**). **d**, Heatmaps of clustered genes based on the signed gene scores from GO enrichment analysis of the core transcriptional regulators. **e**, Schematic of cytosolic redox pathways including KEAP1/NFE2L2 signaling, selenocysteine synthesis, CoQ synthesis, glycolysis, etc. Genes are annotated with modes corresponding to gene effects and scores from batch retest basal and 100 nM RSL3-treated expression (BFP) screens. **f**, Heatmaps of clustered genes based on the signed gene scores from GO enrichment analysis of the core post-translational regulators.

Analysis of enriched sgRNAs using casTLE identified transcriptional and post-translational regulators of FSP1 (**Fig. 3B, C, Table S3**). Overall, the basal and RSL3 screens exhibited a high correlation, indicating that FSP1 regulation is not generally altered in response to GPX4 inhibition (**Extended Data Fig. S2C, D, Table S3**). As expected, *FSP1* was a top hit in the transcriptional screen (BFP), exhibiting strong enrichment in BFP^low^ cells due to disruption of FSP1 expression (**Fig. 3B, Extended Data Fig. S2E, F**). In contrast, *FSP1* sgRNAs were not enriched in the GFP:BFP^high^ or GFP:BFP^low^ cells in the post-translational stability screen (**Fig. 3C, Extended Data Fig. S2E, F**). These results highlight the robust capability of our screening approach to distinguish between transcriptional and post-translational regulators.

Our analyses revealed several interesting functional clusters of *FSP1* transcriptional regulators (i.e., high gene scores in the BFP screen) (**Fig. 3D**). Consistent with prior findings implicating NFE2L2 (also known as NRF2) in FSP1 regulation^17^, *KEAP1* and *NFE2L2* emerged as *FSP1* transcriptional regulators. *KEAP1* disruption increased FSP1 expression, while *NFE2L2* disruption decreased it. Intriguingly, we also identified NFE2L1 as a transcriptional regulator of *FSP1*, suggesting that FSP1 may contribute to the observed ferroptosis suppression by this pathway^28,29^. A cluster of genes associated with selenoprotein biosynthesis and oxidative stress had notable effects on *FSP1* transcription. Although GPX4 is a key ferroptosis resistance factor with an active site selenocysteine, it was not identified in the screen and RSL3 treatment did not induce FSP1 expression (**Fig. 3D, E**). However, depletion of TXNRD1, which also contains an active site selenocysteine, triggered FSP1 upregulation (**Fig. 3D, E**), suggesting a feedback mechanism that responds to certain types of oxidative stress. Another striking cluster influencing *FSP1* transcription included genes involved in acetylation, including the NatB (NAA20, NAA25) and NatC (NAA30, NAA35, NAA38) complexes (**Fig. 3D**). The NatB and NatC complexes are best known for their roles in regulating protein N-terminal acetylation and protein degradation via the N-end rule pathway^30–33^. However, these factors did not exhibit significant scores in the GFP:BFP stability screen (**Fig. 3D**), suggesting that they are either exhibiting unique functions on unknown substrates or regulating *FSP1* transcription indirectly.

Clusters of genes affecting FSP1 stability (i.e., genes enriched in GFP:BFP^high^ or GFP:BFP^low^ cells), were also identified (**Fig. 3F**). A prominent cluster included genes involved in ubiquitination, with the strongest effects observed upon depletion of the E3 ligases UBR4, KCMF1, and RNF126 (**Fig. 3F**). UBR4 and KCMF1 form a large heterodimeric E3 complex that mediates the addition of mixed-linkage ubiquitin chains, promoting the degradation of diverse substrates^33–36^. Meanwhile, RNF126 is implicated in protein quality control pathways, particularly in the degradation of transmembrane proteins that fail to properly integrate into membranes^37^. Our genetic screens suggest that UBR4, KCMF1, and RNF126 contribute to the basal turnover of FSP1.

One of the most striking effects on FSP1 stability was elicited by *RFK* and *FLAD1* sgRNAs which were enriched in GFP:BFP^low^ cells (**Fig. 3C, F**), indicative of a reduction in FSP1 post-translational stability. FAD biosynthesis is a two-step reaction where RFK mediates the first and rate-limiting step by phosphorylating riboflavin (vitamin B2) to create flavin mononucleotide (FMN) which is then adenylated by FLAD1 to form FAD (**Fig. 4A**). Given that FSP1 is an FAD-coordinating protein and an FAD-dependent oxidoreductase, this finding suggests a link between vitamin B2 availability, FAD synthesis, and FSP1 stability.

**Fig. 4.**
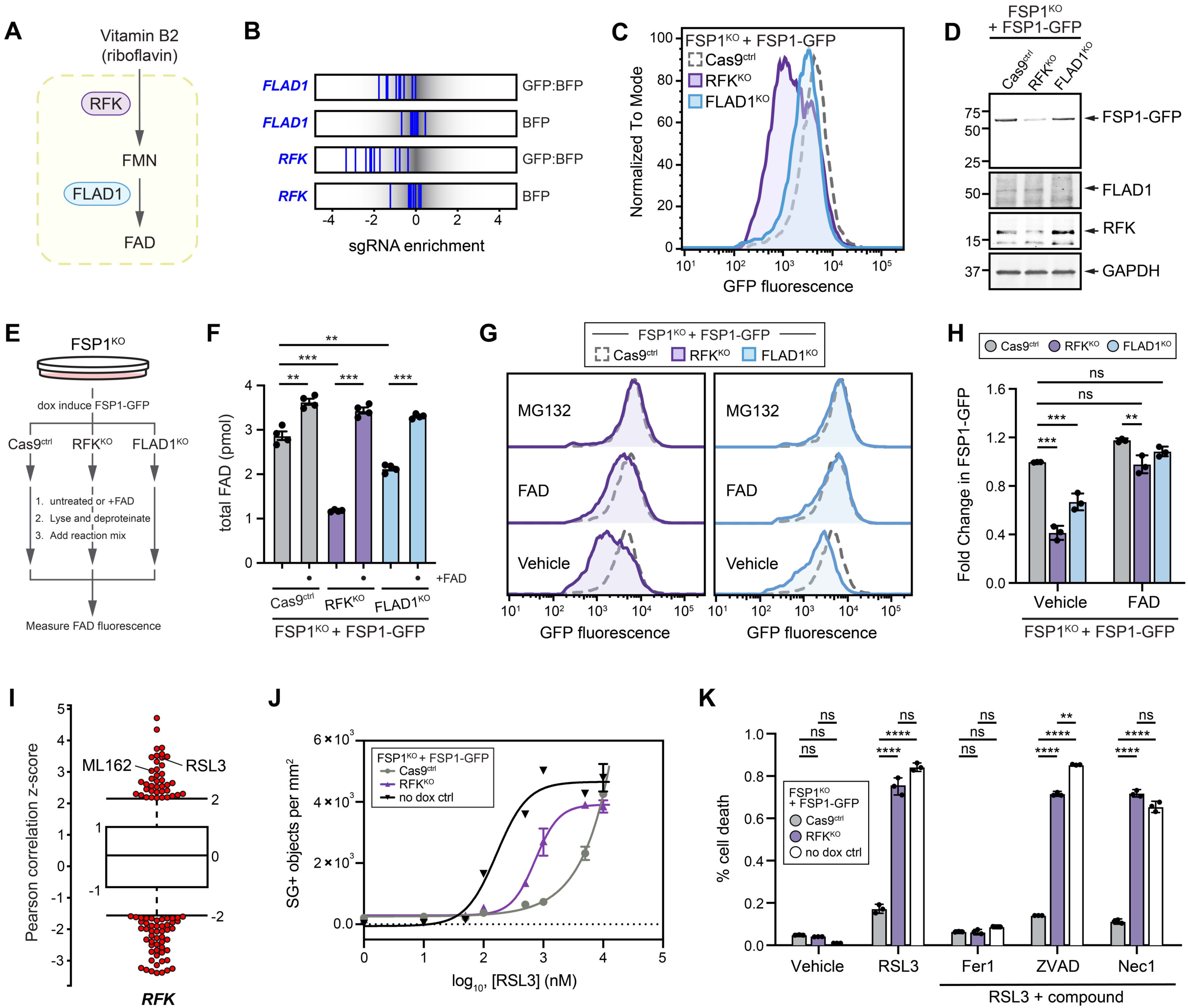
Severe riboflavin deficiency leads to FSP1 protein instability and ferroptosis sensitivity. **a**, Schematic of riboflavin metabolism. **b**, Frequency histograms indicating the distribution of the relative enrichment of *RFK* and *FLAD1* sgRNAs (blue lines) from the batch retest (BFP and GFP:BFP) screens. The gray line indicates the mean of the control sgRNA distribution. **c**, Fluorescence histograms of FSP1^KO^ cells that express GFP-tagged FSP1 in the indicated cell lines. **d**, Immunoblot of lysates from (**c**). **e**, Schematic of FAD detection assay. **f**, FAD quantification of FSP1^KO^ cells expressing FSP1-GFP in the indicated cell lines with and without the presence of 50 µM exogenous FAD for 24 h. **g**, Fluorescence histograms of FSP1^KO^ cells expressing FSP1-GFP in the indicated cell lines treated with 50 µM exogenous FAD or 1 µM MG132. **h**, Quantification of median fluorescence intensity (MFI) change in GFP from (**g**). **i**, RFK expression is positively correlated to ferroptosis inducers, RSL3 and ML162, in cancer cells. Plotted data were mined from the CTRP database, which contains correlation coefficients between gene expression and drug sensitivity for 860 cancer cell lines treated with 481 compounds. **j-k**, Dose response of RSL3-induced cell death (**j**) and percent cell death in the presence of various cell death inhibitors (1 µM RSL3, 2 µM Fer-1, 20 µM ZVAD, and 1 µM Nec-1, **k**) in FSP1^KO^ cells expressing FSP1-GFP in the indicated cell lines treated ± doxycycline for 24 h.

Together, our genetic screens provide insights into FSP1 regulation, effectively stratifying transcriptional and post-translational regulators of FSP1 **(Extended Data Fig. S3A, B)**.

### FAD synthesis promotes FSP1 stability and ferroptosis suppression

Ferroptosis is regulated by several vitamins with diverse mechanisms of action. For example, vitamin E is a well-established suppressor of ferroptosis, functioning as an RTA to inhibit the propagation of lipid peroxidation^1^. Similarly, vitamin K acts as an RTA that is recycled by FSP1^15,16^ and VKORC1L1^38^, further contributing to ferroptosis resistance. Vitamin A also suppresses ferroptosis through multiple mechanisms: its derivatives, retinal and retinol, act as RTAs, while all-trans retinoic acid activates the retinoic acid receptor, which regulates the expression of anti-ferroptotic genes^39^. In contrast, the connection of Vitamin B2 with ferroptosis remains mostly unexplored. The identification of *RFK* and *FLAD1* in our genetic screens suggests that the synthesis of FAD from vitamin B2 promotes FSP1 post-translational stability (**Fig. 3C**). Supporting this hypothesis, multiple sgRNAs targeting *RFK* (9 out of 10 sgRNAs) and *FLAD1* (7 out of 10 sgRNAs) were significantly enriched in GFP:BFP^low^ cells with no enrichment in the BFP screen, identifying these genes as high confidence candidate regulators of FSP1 post-translational stability (**Fig. 4B, Extended Data Fig. S4A, B**).

To evaluate the role of RFK and FLAD1 in FSP1 regulation, we generated *RFK* and *FLAD1* knockout (KO) cells (RFK^KO^ and FLAD1^KO^ cells) in the FSP1^GFP-P2A-BFP^ cell line. Consistent with our genetic screens, loss of RFK and FLAD1 significantly reduced the GFP:BFP ratio and decreased levels of both untagged FSP1 and FSP1-GFP proteins (**Extended Data Fig. S4C-E).** Similarly, loss of RFK and FLAD1 also resulted in lower FSP1-GFP fluorescence in cells expressing ectopic wild-type FSP1-GFP (**Fig. 4C**). Western blot analysis corroborated these findings showing decreased levels of FSP1-GFP protein (**Fig. 4D**). As anticipated, RFK and FLAD1 KO led to reduced cellular FAD levels (**Fig. 4E, F**). Exogenous supplementation of FAD rescued both the diminished FAD levels and restored FSP1-GFP expression to normal levels (**Fig. 4E-H**). These findings indicate that FAD synthesis by RFK and FLAD1 is required for FSP1 post-translational stability, with stronger effects consistently observed following depletion of RFK.

To further explore the connection between RFK and ferroptosis resistance, we analyzed RFK expression and correlations in the Cancer Therapeutics Response Portal^40^. Based on expression and toxicity data from 860 cancer cell lines, it was found that *RFK* expression positively correlates with cancer cell resistance to the GPX4 inhibitors ML162 and RSL3 (**Fig. 4I**). *FLAD1* did not show an appreciable correlation with ferroptosis inducers. These data suggest that the RFK-dependent biosynthesis of FAD from vitamin B2 plays an important role in enabling cancer cells to suppress ferroptosis. To test this hypothesis, we conducted RSL3 dose-response analyses in RFK^KO^ cells. RFK^KO^ cells expressing WT FSP1 showed increased sensitivity to RSL3-induced cell death compared to the control cells (**Fig. 4J**). Moreover, FSEN1 treatment in RFK^KO^ cells had no additive effects (**Extended Data Fig. S4F**). Thus, although loss of RFK likely influences a variety of FAD-dependent cellular processes, these data indicate that the effect of RFK^KO^ on ferroptosis resistance is through its regulation of FSP1. We noted that the RFK^KO^ cells were not as sensitive as the FSP1^KO^ cells (**Fig. 4J**), potentially reflecting some amount of FAD in the serum that cells are grown in. The cell death in the RSL3-treated RFK^KO^ cells was suppressed by the ferroptosis inhibitor and RTA ferrostatin-1 (Fer1), but not the apoptosis inhibitor ZVAD or the necroptosis inhibitor Nec1 (**Fig. 4K**), confirming that loss of RFK sensitizes cells to ferroptotic cell death. Collectively, these results underscore the importance of the vitamin B2-FAD pathway in maintaining FSP1 stability and ferroptosis resistance in cancer.

### FAD binding is required for FSP1 stability

Vitamin B2 is an essential vitamin for maintaining cellular function, and depletion of vitamin B2 can have general effects on cellular viability and proliferation. As an FAD-dependent oxidoreductase, FSP1 relies on FAD binding for function. This suggests that FSP1 instability triggered by disrupting FAD biosynthesis may be direct, reflecting a requirement of FAD binding for FSP1 to maintain its cellular stability. The crystal structure of human FSP1 was recently solved^41–43^, and in the structure the ribityl moiety of FAD is stabilized through interactions with residue D285 of FSP1 (**Fig. 5A**). Previous studies have shown that the D285N mutation abolishes FSP1’s oxidoreductase activity *in vitro* and its ability to promote ferroptosis resistance in cells^19,44^. However, the effects of this mutation on FAD binding and stability have not been explored.

**Fig. 5.**
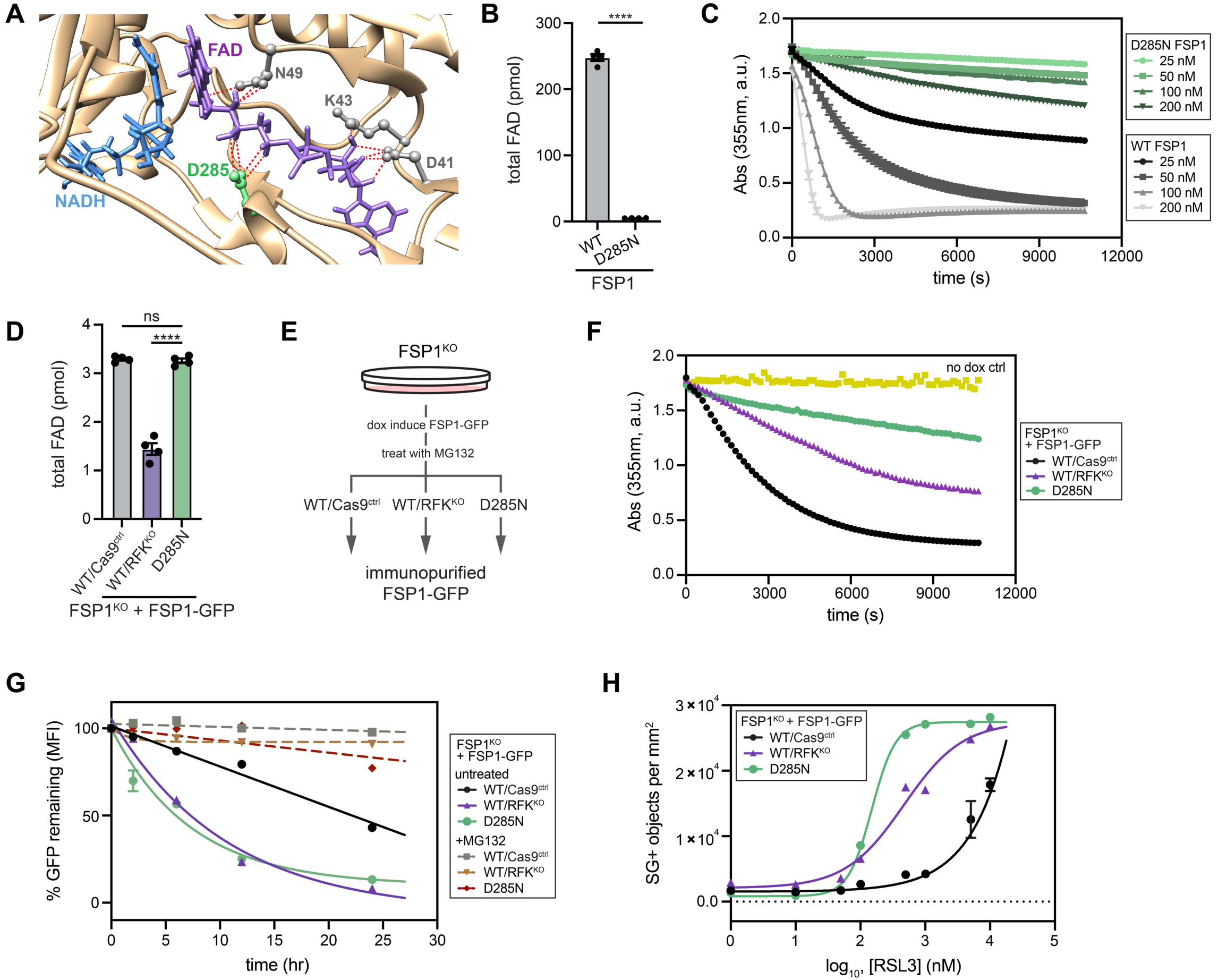
FAD is required for FSP1 function and stability. **a**, Crystal structure of FAD-binding pocket of FSP1 (PDB 8WIK). The cofactors, FAD and NADH, are shown in purple and blue, respectively. FSP1 residues involved in the binding of FAD are shown in stick representation with the D285N interaction highlighted in green. **b**, FAD quantification of recombinant wild-type (WT) and D285N FSP1 proteins. **c**, Oxidation of NADH by recombinant FSP1 variants in the presence of coenzyme Q_1_. **d**, FAD quantification of FSP1^KO^ cells expressing WT or D285N FSP1-GFP in the indicated cell lines. **e**, Purification scheme for the isolation of immunoprecipitated FSP1 from FSP1^KO^ cells expressing FSP1-GFP in the indicated cell lines. **f**, Oxidation of NADH by immunopurified FSP1-GFP variants in the presence of coenzyme Q_1_. **g**, Kinetics of GFP fluorescence decay of FSP1^KO^ cells expressing WT or D285N FSP1-GFP in the indicated cell lines following doxycycline washout. **h**, Dose response of RSL3-induced cell death in FSP1^KO^ cells expressing FSP1-GFP in the indicated cell lines.

To investigate these effects, we purified recombinant WT and D285N mutant FSP1 from bacteria. Notably, the WT FSP1 protein displayed a yellow hue, indicative of bound FAD, whereas the D285N protein was white, consistent with a loss of FAD binding (**Extended Data Fig. S5A**). Measurement of FAD levels confirmed that WT FSP1, but not the mutant D285N FSP1, is bound to FAD (**Fig. 5B**). Additionally, the absorption spectrum of WT FSP1 shows a sharp peak at ∼430 nm, rather than the typical FAD absorbance maxima at ∼375 and ∼450 nm, resembling that of bound 6-hydroxy-FAD^45^ (**Extended Data Fig. S5B**). In contrast to WT FSP1, the D285N mutant produced no noticeable signal (**Extended Data Fig. S5B**), similar to data observed for a D285A mutant^41^. As expected and previously reported^19^, the D285N mutant showed a dramatic reduction in CoQ1 oxidoreductase activity compared to WT FSP1 (**Fig. 5C**). Together, these analyses demonstrate that the D285N mutation disrupts FAD binding and abolishes enzymatic activity.

To examine the importance of FSP1-FAD binding in cells, we analyzed WT FSP1 expressed in RFK^KO^ cells, which have low FAD, and mutant D285N FSP1 expressed in ctrl cells (**Fig. 5D**). In contrast to WT FSP1 from ctrl cells, immunopurified WT FSP1 from RFK^KO^ cells and immunopurified mutant D285N FSP1 from ctrl cells showed reduced CoQ1 oxidoreductase activity (**Fig. 5E, F, Extended Data Fig. S5C**). These data are consistent with our recombinant protein studies where diminished FAD binding or availability hinders FSP1 enzymatic activity. To investigate FSP1 turnover, we utilized a doxycycline (dox)-inducible expression system which encodes an mRNA transcript that lacks a poly-A tail, thereby allowing rapid and selective transcript degradation after dox withdrawal^34,46^. This approach improved dynamic range and enabled analysis of protein turnover with minimal interference from new protein synthesis^34,46^. WT FSP1 was highly stable, with a half-life of approximately 24 hours (**Fig. 5G**). Inhibition of the proteasome with MG132 fully stabilized WT FSP1 (**Fig. 5G**), consistent with its basal turnover being mediated by the ubiquitin-proteasome system. In contrast, both WT FSP1 in RFK^KO^ cells and D285N FSP1 in ctrl cells exhibited significantly reduced stability, with half-lives of approximately 6 hours (**Fig. 5G, Extended Data Fig. S5D, E**). Proteasome inhibition restored stability in both cases (**Fig. 5G**). Finally, we evaluated ferroptosis sensitivity. Unlike WT ctrl cells, RFK^KO^ cells expressing FSP1-GFP or ctrl cells expressing the D285N mutant were more sensitive to ferroptosis (**Fig. 5H**), further underscoring the critical role of FAD binding in FSP1-mediated ferroptosis suppression.

Collectively, our complementary results from recombinant protein studies and cellular analyses demonstrate that disruption of FAD binding—whether by FSP1 mutation or inhibition of FAD synthesis—leads to the generation of an inactive FSP1 protein that is targeted for proteasomal degradation. These findings emphasize the importance of FAD availability and binding for FSP1 stability, function, and ferroptosis resistance.

### RNF8 mediates the turnover of FAD-free FSP1

Our findings indicate that reduced FAD levels or impaired FAD binding leads to FSP1 instability in cells and its degradation by the proteasome. However, the mechanisms underlying the degradation of FAD-free FSP1 remain unclear. Our initial genetic screens identified the E3 ligases UBR4 and KCMF1, which form a heterodimeric complex, suggesting their potential involvement in FSP1 turnover. Consistent with our screen results, depletion of UBR4 or KCMF1 increased the GFP:BFP ratio in FSP1^GFP-P2A-BFP^ cells; however, depletion of UBR4 or KCMF1 did not restore the GFP:BFP ratio FAD-deficient RFK^KO^ cells (**Extended Data Fig. S6A**). These results suggest that while UBR4 and KCMF1 mediate FSP1 turnover under basal conditions, they are not responsible for the degradation of FAD-free FSP1 under conditions of FAD deficiency.

To identify the ubiquitination machinery responsible for the turnover of FAD-free FSP1, we conducted a genetic screen in FSP1^GFP-P2A-BFP^ reporter cells, in which we deleted RFK to disrupt FAD synthesis. We employed a degradation-focused sgRNA library^46^ composed of guides targeting ∼2,000 genes involved in the <u>ub</u>iquitin, <u>a</u>utophagy, and lysosomal (UBAL) degradation pathways and isolated cells using FACS based on the GFP:BFP ratio (**Fig. 6A**). As expected, proteasomal subunits were highly enriched (**Fig. 6B, C, Extended Data Fig. S6B, Table S4**). Additionally, several candidate E3 ligases were identified (**Fig. 6B, C, Extended Data Fig. S6B-D**). Among the top candidates, depletion of RNF8 produced the most significant stabilization of FSP1^GFP-P2A-BFP^ in RFK^KO^ cells (**Extended Data Fig. S6E**). Flow cytometry and western blot analyses confirmed that loss of RNF8 (RNF8^KO^) rescued FSP1-GFP levels in RFK^KO^ cells with no impact to FSP1-GFP levels in the ctrl cells (**Fig. 6D-F, Extended Data Fig. S7A**). Moreover, depletion of RNF8 significantly increased the half-life of ectopically expressed FSP1-GFP in RFK^KO^ cells (**Fig. 6G, Extended Data Fig. S7B**), indicating reduced degradation of FAD-free FSP1. Impaired degradation of FSP1 in the RFK and RNF8 double knockout (RFK/RNF8^DKO^) cells was increased by reintroducing wild-type RNF8, but not a catalytically inactive variant (RNF8^C406S^) (**Fig. 6H**), confirming that E3 ligase activity of RNF8 is required for efficient turnover of FAD-free FSP1. Furthermore, the protein-protein interaction of FAD-free FSP1 with RNF8 in RFK^KO^ cells was confirmed by co-immunoprecipitation, immunoblotting for a C-terminal S-tagged RNF8 and by the reciprocal experiment in which S-tagged RNF8 immunoprecipitants were probed with a FSP1 antibody (**Fig. 6I**). Thus, recognition of FSP1 by RNF8 is only possible when FSP1 is expressed under conditions of FAD deficiency.

**Fig. 6.**
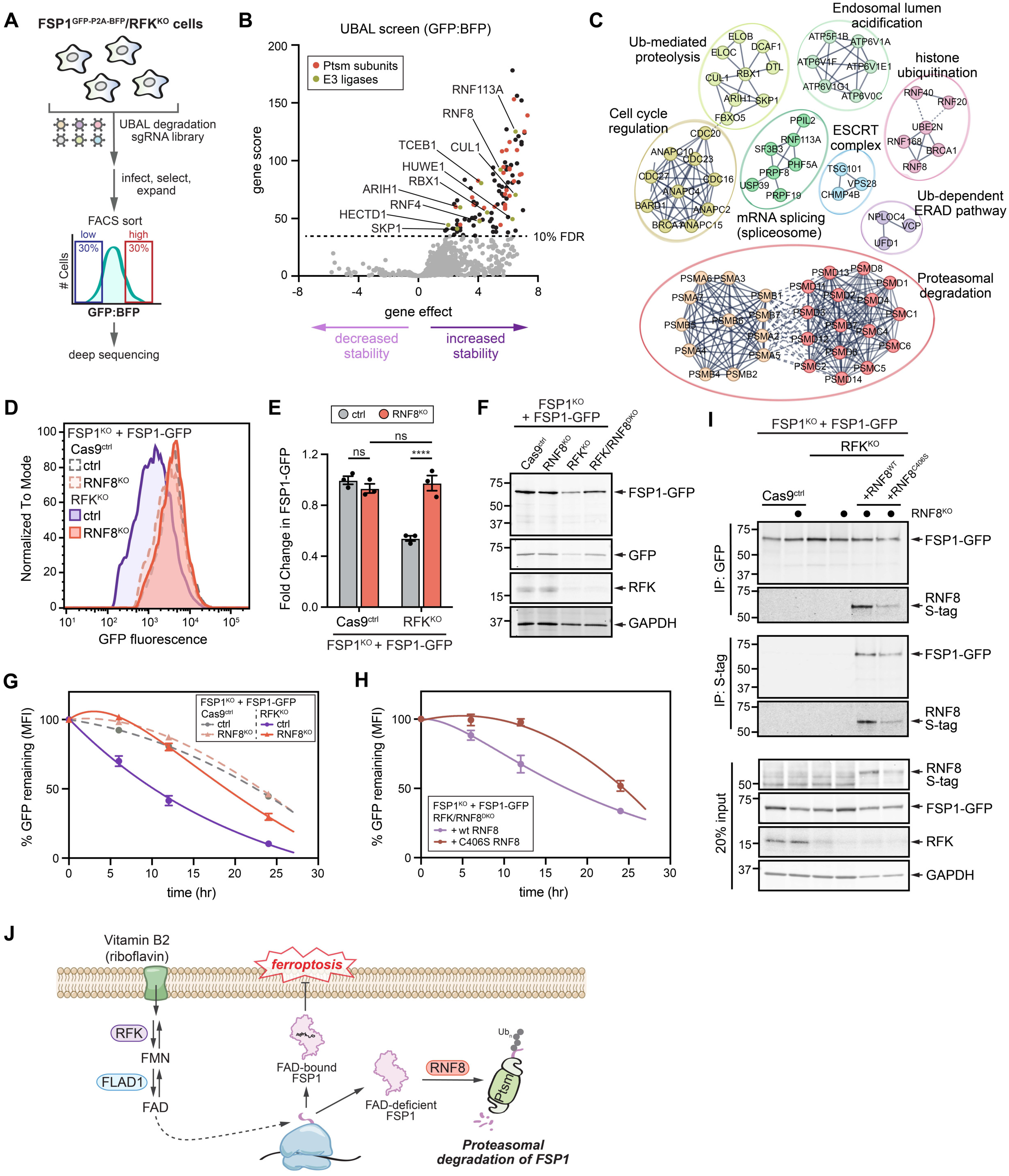
The E3 ligase RNF8 degrades FAD-free FSP1. **a**, Schematic of sensitized UBAL degradation CRISPR-Cas9 screen. **b**, Gene effects and gene scores calculated for individual genes analyzed in the targeted degradation screen. **c**, STRING network map of the identified genes with a 10% FDR cutoff, where each bubble color represents an enriched GO term functional cluster. **d**, Fluorescence histograms of FSP1^KO^ cells expressing FSP1-GFP in the indicated cell lines. **e**, Quantification of median fluorescence intensity (MFI) change in GFP from (**d**). **f**, Immunoblot of lysates from (**d**). **g-h**, Kinetics of GFP fluorescence decay of FSP1^KO^ cells expressing FSP1-GFP in control or RFK^KO^ cells with or without the loss of RNF8 (**g**) or RFK/RNF8^DKO^ (double knock-out) cells expressing the indicated RNF8 variants (**h**) following doxycycline washout. **i**, FSP1-GFP or RNF8 S-tag immunoprecipitation from FSP1^KO^ cells expressing FSP1-GFP in the indicated cell lines and blotting for RNF8 (top) or FSP1 (middle). **j**, Model illustrating the mechanism by which vitamin B2 metabolism and FAD synthesis impacts FSP1 activity and stability to prevent ferroptosis.

To independently validate that RNF8 recognizes FAD-free FSP1, we generated RNF8^KO^ cells in the mutant D285N FSP1 cell line. Loss of RNF8 significantly stabilized FSP1-GFP fluorescence in cells expressing ectopic D285N FSP1-GFP (**Extended Data Fig. S7C, D**). Similarly, depletion of RNF8 increased the half-life of ectopically expressed D285N FSP1-GFP, emphasizing inhibited degradation of FAD-free FSP1 (**Extended Data Fig. S7E, F**).

Together, these findings demonstrate that FAD-free FSP1 is degraded via a ubiquitination pathway involving the E3 ligase RNF8 and the proteasome. This highlights a distinct quality control mechanism for the turnover of FAD-free FSP1 (**Fig. 6J**).

## DISCUSSION

Our genetic screens in our genome-edited FSP1^GFP-P2A-BFP^ reporter cells provide a valuable resource for defining regulators of FSP1 abundance, uncovering many factors and pathways that influence FSP1 expression and post-translational stability. Our findings specifically highlight the pivotal role of vitamin B2 metabolism in FSP1 regulation. Specifically, our results support a model (**Fig. 6J**) in which the conversion of vitamin B2 into FAD facilitates the enzymatic activity and cellular stability of FSP1, shielding it from degradation via a proteasomal pathway involving the E3 ligase RNF8. While several vitamins have been linked to ferroptosis regulation, our findings illuminate a distinct and critical role for vitamin B2 in ferroptosis resistance, that is uniquely tied to the stability and function of FSP1.

Vitamins are well-established players in ferroptosis regulation. Vitamin E, vitamin K, and vitamin A act as RTAs, directly neutralizing lipid peroxyl radicals to prevent ferroptosis^1,15,16,38,38,39^. Additionally, derivatives of vitamin A promote anti-ferroptosis gene expression programs, providing a dual mechanism of protection^39^. Our work expands this framework by uncovering a role for vitamin B2 in ferroptosis, but we demonstrate that vitamin B2 acts through a distinct mechanism in which its derivative, FAD, supports FSP1 stability and function. FAD, which is directly bound to FSP1, is an essential cofactor for its activity as an oxidoreductase, contributing to the reduction of its substrates CoQ10 and vitamin K^13–16,19,41^. Our biochemical and cellular data indicate that FAD binding is required for its enzymatic activity and cellular stability. While we identified both RFK and FLAD1, we consistently observed stronger effects of FSP1 with *RFK* KO compared to *FLAD1* KO. Some flavoproteins can also bind the intermediate FMN, and one possibility is that FSP1 binding to FMN (or similar analogs) could influence its stability. We find that FAD-free FSP1, generated by inhibition of FAD synthesis in cells or disruption of FSP1 FAD binding by mutation, is targeted for degradation by the ubiquitin-proteasome system. This degradation pathway is mediated, in part, by the E3 ligase RNF8, which interacts with FAD-free FSP1 and whose KO stabilizes FSP1 under FAD-deficient conditions.

Binding of the FAD cofactor is important for the cellular stability of some flavoproteins, as the abundance of a subset of flavoproteins decrease in cells grown in the absence of vitamin B2^47^. For example, FAD-free NQO1 or a cancer-associated P187S mutant form of NQO1, which exhibits reduced FAD binding and is degraded by the ubiquitin-proteasome system^47,48^. In the absence of FAD binding, NQO1 is degraded by the quality control E3 ligase C-terminal Hsp70-interacting protein (CHIP)^47^. CHIP was not identified in our genetics screens as a regulator of FSP1 turnover; however, it is potentially notable that RNF8 exhibits conserved structural similarities to CHIP^49^. Although the lack of FAD destabilizes a subset of flavoproteins^47^, our data suggest that the observed ferroptosis sensitizing effects in the RFK^KO^ cells are driven by the reduction in FSP1, as there are no additive effects between FSP1 inhibition and RFK depletion.

Our findings have therapeutic implications since the availability of vitamin B2 and its metabolism influences FSP1 levels and its ability to suppress ferroptosis. The observation that RFK expression correlates with ferroptosis resistance in cancer cells suggests that FAD could be limiting under certain conditions. Targeting this pathway, with therapeutics or diet, could provide a novel strategy to modulate ferroptosis in disease contexts. Moreover, it is also noteworthy that mutations in genes involved in FAD metabolism have been implicated in rare diseases, including mutations in the vitamin B2 transporters (*SLC52A1-3*) and *FLAD1*^50^. Mutations in *RFK* have yet to be identified. Whether FSP1 levels and ferroptosis sensitivity are affected and contribute to the pathogenesis of these diseases are unknown.

Beyond vitamin B2, our genetic screens provide a wealth of additional insights into FSP1 regulation. For instance, we identify known and novel regulators of FSP1 expression, such as NFE2L1 and NFE2L2^17^. We also find that disruptions in oxidative stress and selenoprotein-related genes greatly influence FSP1 expression, raising the possibility that FSP1 is subject to a regulatory feedback mechanism that modulates oxidative stress responses. Our data further validate the regulation of FSP1 stability by the ubiquitin-proteasome system, with UBR4, KCMF1, and RNF126 contributing to its basal degradation. RNF126 has previously been shown to mediate K48-linked polyubiquitination of FSP1 but it was suggested to mediate non-degradative ubiquitination that alters FSP1 localization^51^. Yet, RNF126 is extensively characterized as a protein quality control E3 ligase for integral membrane proteins that fail to insert into the membrane^51^. Our screen results raise the possibility that RNF126 contributes to the turnover of FSP1, in line with its canonical cellular role. Whether it mediates the degradation of mislocalized FSP1 remains to be determined. Although beyond the scope of the current study, understanding the cognate E3 ligases that govern FSP1 degradation is an important goal that may facilitate the development of FSP1 molecular glue degraders. Our findings underscore the complexity of FSP1 regulation and highlight numerous pathways and factors that warrant further investigation to fully characterize their roles in FSP1 regulation and ferroptosis.

In conclusion, our study elucidates a model in which vitamin B2-derived FAD regulates activity and cellular stability, thereby promoting ferroptosis resistance. This discovery highlights a previously unrecognized role for vitamin B2 in ferroptosis and opens new avenues for investigating the interplay between vitamins, FAD, and ferroptosis regulation in physiology and disease.

## AUTHOR INFORMATION

Correspondence and requests for materials should be addressed to J.A.O.

## ACKNOWLEDGEMENTS

This research was supported by a grant from the National Institutes of Health to J.A.O. (R01GM112948) and Bakar Fellows Spark Award to J.A.O. This work was supported in part by funding to J.A.O. as a Chan Zuckerberg Biohub – San Francisco Investigator.

## AUTHOR CONTRIBUTIONS

K.K.D. and J.A.O. conceived of the project, designed the experiments, and wrote the majority of the manuscript. All authors read, edited, and contributed to the manuscript. K.K.D. performed the majority of the experiments. C.A.H. assisted with cell death assays, S.J.T. assisted with chase experiments, C.E.D. purified recombinant FSP1 proteins, and A.J.M. assisted with imaging.

## METHODS

### Cell lines and culture conditions

HEK293T and U-2 OS were all obtained from UC Berkeley Cell Culture Facility. All cell lines were cultured in DMEM medium containing 4.5 g/L glucose, L-glutamine, without sodium pyruvate (Corning, no. 10-013-CMR). All media were supplemented with 10% fetal bovine serum (Gemini Bio Products) and all cells were maintained at 37°C and 5% CO_2_. Penicillin-streptomycin (Life Technologies, no. 15140122) was added to growth media for CRISPR-Cas9 screens and fluorescence-activated cell sorting. All cell lines were tested for mycoplasma.

### Generation of CRISPR-Cas9 genome-edited cell lines

The endogenous, dual-fluorescent FSP1^GFP-P2A-BFP^ knock-in reporter cell line was generated by co-transfection of U-2 OS cells with the donor plasmid pUC57 (described in ‘Plasmids and siRNA’) and px330, a gift from F. Zhang (Addgene plasmid no. 42230), encoding *FSP1* sgRNA guide 1 at a 2:1 w/w ratio using X-tremeGENE HP. 6 h post-transfections, cells were treated and maintained in medium containing 1 µM SCR7 for one week. GFP-P2A-BFP knock-in reporter cells were enriched via six rounds of FACS, and individual clones were isolated using limiting dilution.

For the CRISPR-Cas9 genetic screens, FSP1^GFP-P2A-BFP^ and FSP1^GFP-P2A-BFP^/RFK^KO^ knock-in lines stably expressing Cas9 were generated by transduction with lentiCRISPRv2 hygro virus, a gift from B. Stringer (Addgene plasmid no. 98291), encoding either SAFE-control or *RFK*-targeting sgRNA guides. Cells were selected and maintained in medium containing 375 µg/mL hygromycin. Active Cas9 expression was validated by flow cytometry analysis following infection with a self-cutting mCherry plasmid, which expresses mCherry and an sgRNA targeting the mCherry gene. FLAD1^KO^, UBR4^KO^, KCMF1^KO^, CUL1^KO^, RNF4^KO^, RNF113A^KO^, ARIH1^KO^, and RNF8^KO^ lines were generated by transduction with pMCB320 virus, a gift from M. Bassik (Addgene plasmid no. 89359), encoding the appropriate sgRNA guides. FSP1^GFP-P2A-BFP^ or FSP1^GFP-P2A-BFP^/RFK^KO^ knock-in lines expressing Cas9 were then selected in medium containing 1 µg/mL puromycin and analyzed by flow cytometry.

Lentiviral particles for transduction were generated by co-transfection of lentiCRISPRv2 plasmids with second-generation lentiviral packaging plasmids (pMD2.g and psPAX2) or pMCB320 plasmids with third-generation lentiviral packaging plasmids (pMDLg/pRRE, pRSV-Rev, and pMD2.G) into HEK293T cells using TransIT-LT1 transfection reagent (Mirus) according to manufacturer’s instructions. Lentiviral media was collected 72 h after transfection, passed through a 0.45 µm syringe filter, and used to infect U-2 OS cells in the presence of 8 µg/mL polybrene.

### Generation of doxycycline-inducible cell lines

U-2 OS FSP1^KO^ lines were generated as previously described^13^. Briefly, U-2 OS cells were transfected with px459 (Addgene plasmid no. 48139) encoding *FSP1* sgRNA guide 2, followed by selection in medium containing 1 µg/mL puromycin and isolation of individual clones using cloning rings.

Expression lines were generated by transduction of U-2 OS FSP1^KO^ cells with pLenti CMV rtTA3 Blast (w756-1), a gift from E. Campeau (Addgene plasmid no. 26429), followed by selection in medium containing 10 µg/mL blasticidin. FSP1^KO^ rtTA3-expressing cells were subsequently infected with pMCB497-pTRE-FSP1-GFP-polyA(-)-Blast virus containing either wild-type or D285N FSP1 construct (described in ‘Plasmids and siRNA’). FSP1-GFP expressing cells were enriched by two rounds of fluorescence-activated cell sorting of the GFP-positive populations upon induction with 1 µg/mL doxycycline (dox) for 24 hr.

RFK^KO^ and FLAD1^KO^ lines were generated by transduction with lentiCRISPRv2 hygro (Addgene plasmid no. 98291) virus and selection in medium containing 375 µg/mL hygromycin. RNF8^KO^ cell line was generated by transduction with lentiCRISPRv2 puro (Addgene plasmid no. 98290) virus and selection in medium containing 1 µg/mL puromycin.

RNF8 S-tag rescue lines (described in ‘Plasmids and siRNA’) were generated by transduction with pLenti CMV/TO Zeo DEST (644-1, Addgene plasmid no. 17294) virus containing either wild-type or C406S RNF8 and were selected in medium containing 250 µg/mL zeocin.

All lentiviral particles for transduction were generated by co-transfection with third-generation lentiviral packaging plasmids (pMDLg/pRRE, pRSV-Rev, and pMD2.G) into HEK293T cells, unless otherwise described in ‘Generation of CRISPR-Cas9 genome-edited cell lines.

### Plasmids and siRNA

Cloning of all expression plasmids and the GFP-P2A-BFP donor plasmid was performed using restriction enzyme-independent fragment insertion by megaprimer cloning.

To generate the FSP1 GFP-P2A-BFP knock-in donor plasmid, 800-bp homology arms flanking the FSP1 stop codon were amplified from U2OS genomic DNA and inserted into pUC57 plasmid (a gift from R. Tjian, UC Berkeley). To introduce the GFP-P2A-BFP insert, primers containing 15-bp overlaps with each of the FSP1 homology arms were used to amplify a codon-optimized GFP-P2A-BFP gBlock (Integrated DNA Technologies). This insert was then cloned in frame of the FSP1 stop codon within the pUC57 plasmid containing the FSP1 homology arms. The protospacer adjacent motif site that corresponds to the FSP1 sgRNA guide 1 was subsequently mutated in the donor sequence using site-directed mutagenesis primers without changing the encoded amino acids and to prevent cutting of the integrated donor sequence by Cas9.

Wild-type FSP1 was cloned and inserted in frame 5’ to 3’ to replace the Insig1 gene sequence in pMCB497-pTRE-Insig1-GFP-polyA(-)-Blast plasmid (a gift from R. Kopito, Stanford University). FSP1 D285N was subsequently generated using site-directed mutagenesis. pDONR221 RNF8 (DNASU plasmid no. HsCD00042754) was modified by insertion of a C-terminal S-tag and stop codon at the C terminus of RNF8. RNF8 C406S S-tag was subsequently generated using site-directed mutagenesis. All entry plasmids were cloned into pLenti CMV/TO Zeo DEST by Gateway recombination cloning (ThermoFisher Scientific). For protein expression, wild-type and D285N FSP1 lacking the ATG start codon were inserted into the pET-His6-Tev vector, a gift from S. Gradia (Addgene plasmid no. 29653), C-terminal to the His6-TEV tag.

CRISPR sgRNA sequences targeting *FSP1* were designed using the CRISPR guide design tool by Benchling (https://www.benchling.com). sgRNAs targeting *RFK*, *FLAD1*, *UBR4*, *KCMF1*, *RNF4*, *CUL1*, *RNF113A*, *ARIH1*, *RNF8*, and SAFE-control were selected based on the effective score from the human CRISPR-Cas9 KO libraries used for genetic screening. The oligonucleotide sequences preceding the protospacer motif were: sg*FSP1-*1, 5’-TGAGGCAGTCTCCACCTTGA-3’; sg*FSP1*-2, 5′-GAATCGGGAGCTCTGCACG-3′; sg*RFK*, 5’-GGTCTCAGGTAGCCAACAA-3’; sg*FLAD1*, 5’-GAGTGGAACTGAACAGCT-3’; sg*UBR4*, 5’-GTGCACAGGCCCTTGATT-3’; sg*KCMF1*, 5’-GTGCATCTTGTTATGAAAG-3’; sg*RNF4*, 5’-GGGCTCTGGCAGTGACTAGG-3’; sg*CUL1*, 5’-GGAATTATATAAACGACTTA-3’; sg*RNF113A*, 5’-GCTTTCTCCAGGAAAGGCGG-3’; sg*ARIH1*, 5’-GGAGGAGGATTACCGCTACG-3’; sg*RNF8*, 5’-GACTGTAGGACGAGGATT-3’; sgSAFE control, 5’-AAATTTCATGGGAAAATAG-3’. Guide sequences were cloned into the BbsI restriction site of vector px459 or px330, BsmBI site of lentiCRISPRv2, or between the BstXI and BlpI restriction sites of pMCB320.

The siRNA oligos smart pool targeting FSP1 as well as nontargeting control (siCNTL) were purchased from Horizon Discovery, and cells were transfected with the RNAiMAX reagent (Invitrogen) according to manufacturer’s instructions.

### Genomic DNA Sequencing and Indel Analysis

Primers were designed to flank the either the stop codon or predicted Cas9 cut site, with the forward primer ∼250bp upstream of the cut site.

For the genomic DNA extraction, cells were washed twice and scraped off plates in cold DPBS and pelleted at 500*g* for 5 min. Genomic DNA was extracted from cell pellets and purified using the Qiagen Blood Mini Kit (Qiagen, no. 51104) according to manufacturer’s protocol.

PCR was performed on an Applied Biosystems Thermal Cycler using Q5 High-Fidelity 2X Master Mix (New England Biolabs, no. M0492S). TIDE oligonucleotide sequences are as listed:

Fwd-*RNF8*-Exon2, 5’-CTACTTGGTGGTTCTCAAGACAGC-3’; Rev-*RNF8*-Exon2, 5’-GTCTCTGGGAGCTTCACCTCA-3’.

Amplicons were separated on 2% agarose-TAE gels and purified using the QIAquick Gel Extraction Kit (Qiagen, no. 28706). Amplicons were sequenced at QuintaraBio, and sequences were assessed for indels using TIDE analysis tool (http://shinyapps.datacurators.nl/tide/).

### Chemicals and reagents

We purchased 1S,3R-RSL3 (no. 19288), FSEN1 (no. 38025), Ferrostatin-1 (no. 17729), idebenone (no. 15475), DFO (no. 14595), ZVAD(OMe)-FMK (no. 27421), and Necrostatin-1 (no. 11658) from Cayman Chemical. 3-Methyladenine (no. 189490), Bafilomycin A1, *Streptomyces griseus* (no. 196000), Bovine Serum Albumin (no. A8806), Oleic acid (no. O1383), Trichloroacetic acid (no. T6399), Potassium hydroxide (no. 221473), Flavin adenine dinucleotide disodium salt hydrate (no. F6625), and Coenzyme Q1 (no. C7956) were all purchased from Sigma-Aldrich. Blasticidin S HCl (no. A1113903), Puromycin (no. A1113803), Hygromycin B (no. 10687010), Zeocin (no. R25001), BODIPY 581/591 C11 (no. D3861), CellMask DeepRed Plasma Membrane Stain (no. C10046) and SYTOX Green Dead Cell Stain (no. 34860) were all purchased from ThermoFisher Scientific. Lipi-Blue (no. LD01) was purchased from Dojindo Laboratories. SCR7 (no. S7742), CB5083 (no. S8101), MLN7243 (no. S8341), MG132 (no. S2619) were purchased from Selleck Chemicals. NADH (no. 481913), and Doxycycline Hyclate (no. D9891) were purchased from Millipore Sigma. Transfection reagents used in this study include: Lipofectamine RNAiMAX Transfection Reagent (no. 13778150, ThermoFisher Scientific), Polybrene (no. 107689, Millipore Sigma), X-tremeGENE HP DNA Transfection Reagent (no. 6366244001, Roche), and TransIT-LT1 Transfection Reagent (no. MIR2300, Mirus).

### Western blotting

Cells were washed twice with phosphate-buffered saline (PBS), lysed in 1% SDS, sonicated for 20 s and incubated for 5 min at 100°C. The cell lysate was then centrifuged for 10 min at 15,000*g* to remove any cell debris. Protein concentrations were determined using the bicinchoninic acid (BCA) protein assay kit (ThermoFisher Scientific), and equal amounts of protein by weight were combined with Laemmli buffer, boiled for 5 min at 100°C, separated on 4–20% polyacrylamide gradient gels (Bio-Rad Laboratories) and transferred onto nitrocellulose membranes (Bio-Rad Laboratories). Membranes were blocked in PBS with 0.1% Tween-20 (PBSt) containing 5% (w/v) dried milk for 60 min. Membranes were washed twice with water and incubated overnight at 4°C in Tris-buffered saline with with 0.1% Tween-20 (TBSt) containing 3% bovine serum albumin (BSA) (Sigma-Aldrich) and primary antibodies. After washing with PBSt, membranes were incubated at room temperature for 60 min in PBSt containing fluorescent secondary antibodies. Immunoblots were imaged on a LI-COR imager (LI-COR Biosciences).

The following blotting reagents and antibodies were used: anti-FSP1 (Santa Cruz, no. sc377120), anti-GFP (Sigma-Aldrich, no. SAB1305545), GAPDH (Cell Signaling, no. D4C6R), anti-PLIN2 (Abcepta, no. AP5118c), anti-Derlin2 (a kind gift from Y. Ye, NIH), anti-RFK (Proteintech, no. 15813-1-AP), anti-FLAD1 (Santa Cruz, no. sc376819), and anti-S-Tag (Novus Biologicals, no. NB600-511).

### Flow cytometry

For flow cytometry analysis, U-2 OS cells were seeded in 6-well plates. If subjected to treatments, FSP1^GFP-P2A-BFP^ cells were treated the next day with 5 mM 3-MA, 250 nM Baf-1A, 5 µM CB5083, 10 µM MLN7243, and 10 µM MG132 for 6 h and/or varying concentrations of RSL3 for 24 h, while FSP1^KO^ cells expressing an inducible, GFP-tagged FSP1 were treated with 1 µg/mL doxycycline and/or 1 µM MG132 or 50 µM FAD for 24 h. Cells were dissociated from plates using TrypLE Express (Gibco) and resuspended in DMEM containing 10% FBS. Cells were pelleted by centrifugation at 500*g* for 5 min, resuspended in DPBS, and placed on ice.

For all flow cytometry assays, fluorescence was analyzed using an LSR Fortessa (BD Biosciences) collecting at least 20,000 events per sample. The following filter sets were used: FITC (GFP, BODIPY C11^ox^), Pacific Blue (BFP), and Texas-Red (mCherry, BODIPY C11^non-ox^). Data were analyzed using FlowJo version 10.10.0 (FlowJo, LLC).

### Fluorescence microscopy

For super-resolution microscopy of live cells, U-2 OS FSP1^GFP-P2A-BFP^ knock-in cells were seeded in 35 mm imaging dishes (CellVis, D35-14-1.5-N) 48 h before the start of imaging. The next day, cells were incubated with 200 µM oleate for 24 h. Lipid droplets were stained with 500 nM Lipi-Blue for 30 min and the plasma membrane was stained with 5 µg/mL CellMask Deep Red for 10 min prior to imaging. Lattice-SIM images were acquired on a Zeiss Elyra7 super-resolution fluorescence microscope, equipped with dual sCMOS PCO Edge 4.2 cameras for simultaneous two channel acquisition, with a 63x/1.4 oil objective. For each focal plane, 13 phase images were acquired. Lattice-SIM reconstruction was performed with the SIM processing Tool of the ZEN 3.0 SR (Black, v16) software.

### Lipid droplet fractionation

Two 500-cm^2^ plates of U-2 OS FSP1^GFP-P2A-BFP^ knock-in cells were incubated in 200 µM oleate-BSA complex for 24 h. Cells were collected by scraping into PBS and centrifuged for 10 min at 500*g*. Cell pellets were resuspended in cold hypotonic lysis medium (HLM, 20 mM Tris-HCl pH 7.4 and 1 mM EDTA) supplemented with 1× cOmplete, Mini, EDTA-free Protease Inhibitor Cocktail (Sigma-Aldrich), incubated on ice for 10 min, lysed using 20 passes via a syringe affixed to a 23g needle and centrifuged at 1,000*g* for 10 min. The supernatant was subsequently transferred to Ultra-Clear ultracentrifuge tubes (Beckman-Coulter), diluted with 60% sucrose in HLM to a final concentration of 20% sucrose in HLM, and overlaid by 4 mL of 5% sucrose in HLM buffer followed by 4 mL of HLM buffer. Overlaid samples were centrifuged for 30 min at 15,000*g* in an ultracentrifuge using a SW41 swinging bucket rotor. Buoyant fractions were collected using a tube slicer (Beckman-Coulter), additional fractions were pipetted from the top of the sucrose gradient in 1-mL increments, and pellets were resuspended in 1 mL HLM. 100 µL of 10% SDS was added to each fraction, yielding a final concentration of 1% SDS. Samples were then sonicated for 15 s and incubated for 10 min at 65°C. Buoyant fractions were incubated at 37°C for 1 h and sonicated every 20 min, followed by a final incubation at 65°C for 10 min.

### Cell viability and death analysis

Cells were plated in triplicate at a density of 2,000 cells per well in black 96-well plates (Corning, no. 3904) 48 h before the start of imaging. To induce expression of FSP1, cells were treated with 1 µg/mL doxycycline the next day. After 48 h, the medium was replaced with fresh medium containing 30 nM SYTOX Green Dead Cell Stain, doxycycline (if needed) and the indicated drugs at different doses. The plates were immediately transferred to an IncuCyte S3 Live-Cell imaging system (Sartorius) enclosed in an incubator set to 37°C and 5% CO_2_. Three images per well were captured in the green, red, and phase channels every 3 h over a 24 h period. The ratio of SYTOX Green-positive objects (dead cells) to phase objects (total cells) or mCherry-positive objects (live cells) was quantified using the Sartorius IncuCyte 2020A image analysis software. For each treatment condition, the SYTOX-to-mCherry-object ratio (wild-type and FSP1^GFP-P2A-BFP^ lines) or SYTOX-to-phase-object ratio (FSP1-GFP expression lines) was plotted against the 24 h imaging interval, the average AUC (area under the curve) was plotted as a function of drug concentration (for example, RSL3) using GraphPad Prism v10. To compare cell death between FSP1-GFP expression lines (wild-type, RFK^KO^, FLAD1^KO^, and no dox control), the SYTOX-object counts of 24 h timepoint from 1 µM RSL3 treatment were normalized by the maximum value for each cell line (i.e. the phase-object counts of 0 h timepoint).

### Genome-wide U-2 OS FSP1^GFP-P2A-BFP^ CRISPR-Cas9 screen

Genome-wide CRISPR-Cas9 screens were performed using the Bassik Human CRISPR Knockout Library (Addgene pooled libraries nos. 101926-101934). The library consists of nine sub-libraries, comprising a total of 225,171 elements, including 212,821 sgRNAs targeting 20,549 genes (∼10 sgRNAs per gene) and 12,350 negative-control sgRNAs. To generate lentiviral particles, each sub-library was co-transfected with third-generation lentiviral packaging plasmids into HEK293T cells. Media containing lentivirus was collected 48 and 72 h after transfection, combined, and filtered. U-2 OS FSP1^GFP-P2A-BFP^ cells stably expressing Cas9 were transduced with lentiviral packaged sub-libraries (one sub-library at a time) in 8 µg/mL polybrene. After 72 h of growth, infected cells were selected in media containing 1 µg/mL puromycin until over 90% of cells were mCherry positive (via flow cytometry). Cells were then recovered for 3-5 days in media lacking puromycin and frozen in liquid nitrogen.

For the screen, library infected cells were thawed (one sub-library at a time) and expanded at 1000x coverage (1,000 cells per library element). On the day of the sort, cells were dissociated using 0.25% Trypsin-EDTA (Gibco), collected by centrifugation at 400*g* for 5 min, and washed 1x with DPBS. Cells were resuspended in phenol red free media (HyClone, no. 16777-406) supplemented with 3% FBS and 1% BSA (fatty acid free) and kept on ice until FACS.

Cells were sorted on a BD Aria Fusion equipped with 4 Lasers (488nm, 405nm, 561nm, and 640nm). sgRNA-expressing, mCherry+ cells were then sorted to the brightest and dimmest 30% GFP+ populations into 15 mL conicals containing DMEM with 4.5 g/l glucose and L-glutamine supplemented with 10% FBS. For each sub-library sort, at least 1000x as many cells as guides were collected. Sorted cells were collected by centrifugation at 1,000*g* for 10 min, washed 1x with DPBS, and pellets frozen at -80°C until genomic DNA extractions. PCR amplicon preparation and deep sequencing were performed according to Mathiowetz et al.^52^.

### Secondary pooled screens using a custom sgRNA library

The custom sgRNA batch retest library contains 9,350 elements with 7,350 sgRNAs targeting 735 genes (∼10 sgRNAs per gene) and 2,000 negative control sgRNAs. Guide sequences were from the Bassik Human CRISPR Knockout Library, and the library construction protocol was previously described^22,23^.

U-2 OS FSP1^GFP-P2A-BFP^ cells stably expressing Cas9 were transduced with the packaged library as described above. For each screen, cells were thawed and expanded at >1000x coverage. For all screens, cells were seeded into 500-cm^2^ plates at 1,000-fold library coverage. For inducing lipid peroxidation, cells were treated the following day with 100 nM RSL3 for 24 h. Cells were screened by FACS as described above with minor modifications: sgRNA-expressing, mCherry+ cells were sorted as follows: GFP screen (brightest 30% GFP+ and dimmest 30% GFP+), BFP screens (brightest 30% BFP+ and dimmest 30% BFP+), GFP:BFP screens (brightest 30% GFP:BFP+ ratio and dimmest 30% GFP:BFP+ ratio).

### Sensitized degradation screen with U-2 OS FSP1^GFP-P2A-BFP^/RFK^KO^ reporter cell line

The custom UBAL degradation library contains 20,710 elements with 18,710 sgRNAs targeting 1,871 genes (∼10 sgRNAs per gene) and 2,000 negative control sgRNAs as previously described^46^.

U-2 OS FSP1^GFP-P2A-BFP^/RFK^KO^ cells stably expressing Cas9 were transduced with the packaged library as described above. Cells were thawed and maintained at >1000x coverage. For all screens, cells were seeded into 500-cm^2^ plates at 1,000-fold library coverage. sgRNA-expressing, mCherry+ cells were then sorted to the brightest and dimmest 30% GFP:BFP+ ratio by FACS.

### CRISPR-Cas9 screen data analysis

Sequence reads were aligned to the sgRNA reference library using Bowtie 2 software. For each gene, a gene effect and score (likely maximum effect size and score) and *p*-values were calculated using the Cas9 high-Throughput maximum Likelihood Estimator (casTLE) statistical framework as previously described^22,23^.

Functional interactions and protein-protein interactions for high confidence candidate regulators were identified using the GO enrichment analysis and STRING interaction network database.

### Flavin adenine dinucleotide (FAD) measurements

1 x 10^6^ U-2 OS FSP1^KO^ cells expressing the indicated FSP1-GFP construct and/or sgRNA were seeded into 10-cm^2^ plates. The following day, cells were treated with medium containing 1 µg/mL doxycycline for 24 h or 50 µM exogenous FAD when necessary. Cells were washed twice with cold PBS and collected by scraping. For recombinant FSP1 proteins, 10 µg of wild-type or D285N variants were resuspended in 100 µL cold FAD Assay buffer. All samples were prepared for FAD fluorescence measurements using the Flavin Adenine Dinucleotide (FAD) Assay Kit (Abcam) according to the manufacturer’s protocol. FAD concentrations were calculated using a standard curve and normalized to the total cell lysate (500 µg) or total protein level (10 µg) in each sample. Four biological replicates were performed for each condition.

### Protein purification

Expression vectors were transformed into LOBSTR-BL21 (DE3) competent cells (Kerafast, Cat# EC1002) and LB broth was inoculated for overnight growth at 37°C. The following day, the cultures were diluted 1:100 into 1L LB broth, allowed to grow at 37°C until the cultures reached an OD_600_ of 0.6 and induced with 0.5 mM isopropyl b-D-1-thiogalactopyranoside (IPTG) overnight at 20°C. Bacterial pellets were harvested, flash frozen, and resuspended in lysis buffer A (50 mM KH_2_PO_4_ pH 8.0, 300 mM KCl, 10% glycerol, 30 mM imidazole, 1 mg/mL lysozyme, and 1 mM PMSF). The resuspended cells were lysed by sonication and centrifuged for at 50,000*g* for 30 min at 4°C.

All subsequent purification steps were carried out at 4°C. The supernatant was passed through an Econo-Column Chromatography Column (Bio-Rad, no. 7371512) packed and equilibrated with 1 mL HisPur Ni-NTA agarose resin (ThermoFisher Scientific, no.88221), and washed 4x with EQ buffer (50 mM KH_2_PO_4_ pH 8.0, 300 mM KCl, 10% glycerol, 30 mM imidazole). Bound proteins were eluted using EQ buffer containing 250 mM imidazole and buffer exchanged into SEC buffer A (50 mM HEPES pH 8.0, 100 mM KCl, 2 mM DTT). The protein sample was then further purified by gel filtration through a HiLoad 16/600 Superdex 75 Pg size exclusion chromatography column in SEC buffer using a GE Akta Pure FPLC.

Minor modifications were made for the purification of D285N FSP1. Bacterial pellets were resuspended in lysis buffer B (50 mM KH_2_PO_4_ pH 8.0, 100 mM NaCl, 100 mM KCl, 10 mM MgCl_2_, 1 mg/mL lysozyme, 1 mM PMSF, 30 mM imidazole, and 10% glycerol), washed 4x with EQ buffer B (50 mM KH_2_PO_4_ pH 8.0, 100 mM NaCl, 100 mM KCl, 10 mM MgCl2, 30 mM imidazole, and 10% glycerol), and eluted with EQ buffer B containing 250 mM imidazole. The eluate was buffer exchanged into SEC buffer B (50 mM HEPES pH 8.0, 100 mM KCl, 2 mM DTT, and 5% glycerol) and purified as previously described. Combined fractions were concentrated and snap-frozen using liquid N_2_. Protein concentration was determined using BCA protein assays.

### Immunoprecipitations

U-2 OS FSP1^KO^ cells expressing the indicated FSP1-GFP construct and/or sgRNA were seeded into 10-cm^2^ plates. The following day, cells were treated with medium containing 1 µg/mL doxycycline and 1 µM MG132 for 24 h. Cells were washed twice with cold PBS, collected by scraping, and spun down at 1,000*g* for 10 mins. Cell pellets were resuspended in 100 µL lysis buffer C (50 mM Tris-HCl pH 7.5, 150 mM NaCl, 0.5 mM EDTA, 0.5% IGEPAL 630, and 2 mM *N*-ethylmaleimide supplemented with 1× cOmplete, Mini, EDTA-free Protease Inhibitor Cocktail. Lysates were sonicated for 30 s and cleared by centrifuging at 20,000*g* for 15 min. Protein concentrations were determined using BCA protein assay.

For immunopurification of FSP1-GFP proteins, 5 mg of pre-cleared lysates were incubated with 50 µL of ChromoTek GFP-Trap Magnetic Agarose (Proteintech, no. gtma) for 1 h at 4°C with end-over-end rotation. Beads were washed 3 times with PBS and 5 µL of bound proteins were eluted by adding 2x Laemmli buffer and incubating at 100°C for 5 min. Immunopurified FSP1-GFP concentrations were calculated using a standard curve of recombinant FSP1 and normalized via densitometry using the anti-FSP1 antibody.

For immunoprecipitation of FSP1 and RNF8 interactions, 1 mg of pre-cleared lysates were incubated with 10 µL of ChromoTek GFP-Trap Magnetic Agarose or 10 µL of S-protein Agarose (Millipore Sigma, no. 69704) for 1 h at 4°C with end-over-end rotation. Beads were washed 3 times with lysis buffer C and bound proteins were eluted by adding 2x Laemmli buffer and incubating at 100°C for 5 min.

### Protein activity assays

To measure NADH oxidation kinetics, 2.5 pmol of immunopurified FSP1-GFP isolated from indicated cell lines or recombinant wild-type or D285N FSP1 at indicated concentrations were combined with 500 µM NADH and 400 µM coenzyme Q1 in a total volume of 200 µL PBS. A reduction in absorbance at 340 nm, corresponding to NADH oxidation, was determined over the course of 3 h at 37°C. All measurements were taken using a Tecan Spark Multimode Microplate Reader (Tecan).

### FAD absorption spectroscopy

To assess the presence of protein-bound FAD, 100 µg of recombinant wild-type or D285N FSP1 proteins or 10 µg of free FAD control were resuspended in a total volume of 200 µL PBS. Absorption spectra were acquired using a Tecan Spark Multimode Microplate Reader (Tecan). All spectra were acquired over a range of 350 to 500 nm with a wavelength step size of 2 nm, manual gain of 100, and 30 flashes at 25°C. The relative absorption measured was normalized by setting the maximum value to 1 of each given sample.

### Transcriptional shutoff “chase” or protein turnover assays

U-2 OS FSP1^KO^ cells expressing the indicated FSP1-GFP construct and/or sgRNA were seeded into 6-well plates. The following day, cells were treated with medium containing 1 µg/mL doxycycline for 12 h to induce expression of FSP1-GFP. Cells were then washed 1x with DPBS and replenished with complete medium to initiate doxycycline washout and measure reporter protein turnover at the indicated times. MG132 controls were treated with 1 µM MG132 during both doxycycline induction and washout.

For flow cytometry analysis, the median fluorescence intensity (MFI) was calculated using FlowJo v10. Cells without dox were run as controls for background fluorescence, and the average MFI for the “no dox” controls was subtracted from the MFI of the dox positive samples at all time points. GFP percentage remaining was calculated as MFI at *t* = *X* divided by the MFI at *t* = 0 h. One-phase decay curves were drawn using GraphPad Prism v10 using the default settings.

### Statistical analysis and reproducibility

All figures, including western blots, dose-response curves, protein turnover assays and panels are representative of two biological replicates unless stated otherwise. For comparison across multiple experimental groups, *p*-values were calculated using two-way ANOVA, and adjusted using Bonferroni correction for multiple comparisons (***P < 0.001, **P < 0.002, *P < 0.033).

### Reporting summary

Further information on research design is available in the Nature Research Reporting Summary linked to this article.

## Materials availability

All unique/stable reagents generated in this study are available from the <u>lead contact</u> with a completed Materials Transfer Agreement.

## Data availability

All data that support the conclusions in this manuscript are available from the corresponding author upon reasonable request. Raw data for Fig. 2b can be accessed in Supplementary Dataset 1. Raw data for Fig. 2e can be accessed in Supplementary Dataset 2. Raw data for Fig. 3b and 3c can be accessed in Supplementary Dataset 3. Raw data for Fig. 6b can be accessed in Supplementary Dataset 4. Raw data for Fig. 4i is publicly available from the CTRP v2 database (https://portals.broadinstitute.org/ctrp.v2.1/.)

## EXTENDED DATA FIGURE LEGENDS

**Fig. S1 |Generation and characterization of a genome-edited FSP1 reporter cell line. a**, Fluorescence histograms of wild-type and genomic FSP1^GFP-P2A-BFP^ knock-in cells. **b**, Genomic sequencing of the *FSP1* gene in control and FSP1^GFP-P2A-BFP^ knock-in cells. **c**, Dose response of RSL3-induced cell death in FSP1^GFP-P2A-BFP^ cells with various cell death inhibitors (2 µM Fer-1, 100 µM DFO, 10 µM idebenone, 20 µM ZVAD, and 1 µM Nec-1). **d-f**, Fluorescence histograms of FSP1^GFP-P2A-BFP^ cells treated with proteolysis inhibitors for 6 h. **f**, Quantification of median fluorescence intensity (MFI) change in GFP:BFP ratio from (**e**).

**Fig. S2 | Genetic screens for transcriptional and post-translational FSP1 regulators. a**, Distribution of counts across all sgRNA elements cloned into the custom batch retest sgRNA library. **b**, Dose response of RSL3-induced cell death in FSP1^GFP-P2A-BFP^ cells labeled with the lipid peroxidation sensor, BODIPY 581/591 C11. Green and red fluorescence intensity was analyzed by flow cytometry. **c-d**, Scatter plot of signed gene scores for individual genes from batch retest screen of untreated (x-axis) versus RSL3-treated (y-axis) duplicates for BFP sort (**c**) or GFP:BFP sort (**d**). **e**, Cloud plots indicating count numbers corresponding to *FSP1* (color scale) and control (gray scale) sgRNAs. **f**, Frequency histograms indicating the distribution of the relative enrichment for *FSP1* sgRNAs (blue lines). The gray line indicates the mean of the control sgRNA distribution.

**Fig. S3 | Heatmap of top transcriptional and post-translational FSP1 regulators. a-b**, Heatmaps displaying the signed gene scores of the top 40 enriched (left) and depleted (right) genes from the BFP (**a**) and GFP:BFP (**b**) batch retest screens.

**Fig. S4 | Impairments of riboflavin metabolism result in FSP1 protein instability and ferroptosis sensitivity. a-b**, Cloud plots indicating count numbers corresponding to *RFK* (**a**) or *FLAD1* (**b**) (color scale) and control (gray scale) sgRNAs. **c-d**, Fluorescence histograms of FSP1^GFP-P2A-BFP^ cells in the indicated cell lines showing changes in GFP and BFP fluorescence (**c**) or GFP:BFP ratio (**d**). **e**, Immunoblot of FSP1^GFP-P2A-BFP^ lysates from (**c-d**). **f**, Dose response of RSL3-induced cell death in FSP1^KO^ cells expressing FSP1-GFP in the indicated cell lines treated with 2 µM FSEN1.

**Fig. S5 FAD is required to prevent loss of function and protein instability. a**, Characterization of recombinant wild-type (WT) and D285N FSP1 proteins: color difference (left) and Coomassie gel (right). **b**, UV-visible spectrum of WT and D285N FSP1 proteins along with free, unbound FAD control (in yellow). **c**, Quantitative immunoblot analysis to determine the concentration of immunopurified FSP1-GFP from the indicated cell lines using recombinant FSP1 protein (inset western blot). **d**-**e**, Immunoblot of lysates (**d**) and fluorescence histograms of doxycycline washout after 12 h (**e**) from FSP1^KO^ cells expressing FSP1-GFP in the indicated cell lines.

**Fig. S6 Sensitized CRISPR-Cas9 screen identifies degradation machinery of destabilized FSP1. a**, Fluorescence histograms of control (left) or RFK^KO^ (right) FSP1^GFP-P2A-BFP^ reporter cells in the indicated cell lines. **b**, Scatter plot of signed gene scores for individual genes from targeted degradation screen of bio rep 1 (x-axis) versus bio rep 2 (y-axis). **c**, Cloud plots indicating count numbers corresponding to various E3 ligases (color scale) and control (gray scale) sgRNAs. **d**, Frequency histograms indicating the distribution of the relative enrichment for sgRNAs (red lines) of various E3 ligases. The gray line indicates the mean of the control sgRNA distribution. **e**, Fluorescence histograms of control (left) or RFK^KO^ (right) FSP1^GFP-P2A-BFP^ cells in the indicated cell lines.

**Fig. S7 RNF8 targets FAD-free FSP1 for degradation. a**, Genomic sequencing of the *RNF8* gene in RFK^KO^ and RFK^KO^/RNF8^DKO^ cells. **b**, Fluorescence histograms of doxycycline washout after 12 h from FSP1^KO^ cells expressing FSP1-GFP in the indicated cell lines. **c**, Fluorescence histograms of FSP1^KO^ cells expressing D285N FSP1-GFP in the indicated cell lines. **d**, Quantification of median fluorescence intensity (MFI) change in GFP from (**c**). **e**, Kinetics of GFP fluorescence decay of FSP1^KO^ cells expressing D285N FSP1-GFP in control or RNF8^KO^ cells following doxycycline washout. **f**, Fluorescence histograms of doxycycline washout after 12 h from FSP1^KO^ cells expressing D285N FSP1-GFP in the indicated cell lines.

## Supplementary information

Supplementary Information (AI file)

Supplementary Figs.1-5

Supplementary Dataset 1 – Genome-wide CRISPR screen results

Supplementary Dataset 2 – GFP sorted batch retest CRISPR screen results

Supplementary Dataset 3 – BFP or GFP:BFP sorted batch retest CRISPR screen results

Supplementary Dataset 4 – UBAL degradation CRISPR screen results

