## Supplementary figures and images for "Vitamin B2 metabolism promotes FSP1 stability to prevent ferroptosis"

### Supplemental Figures

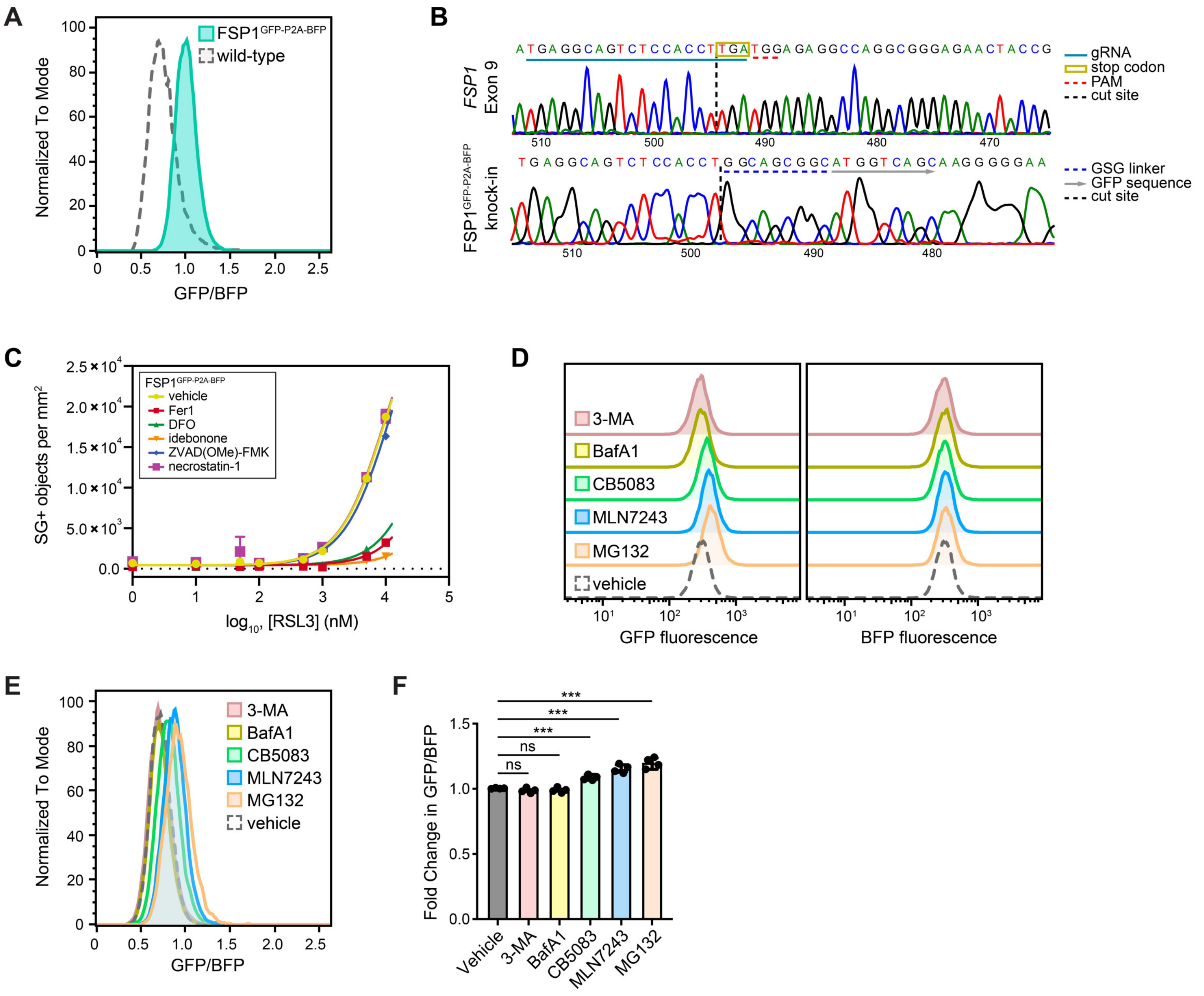

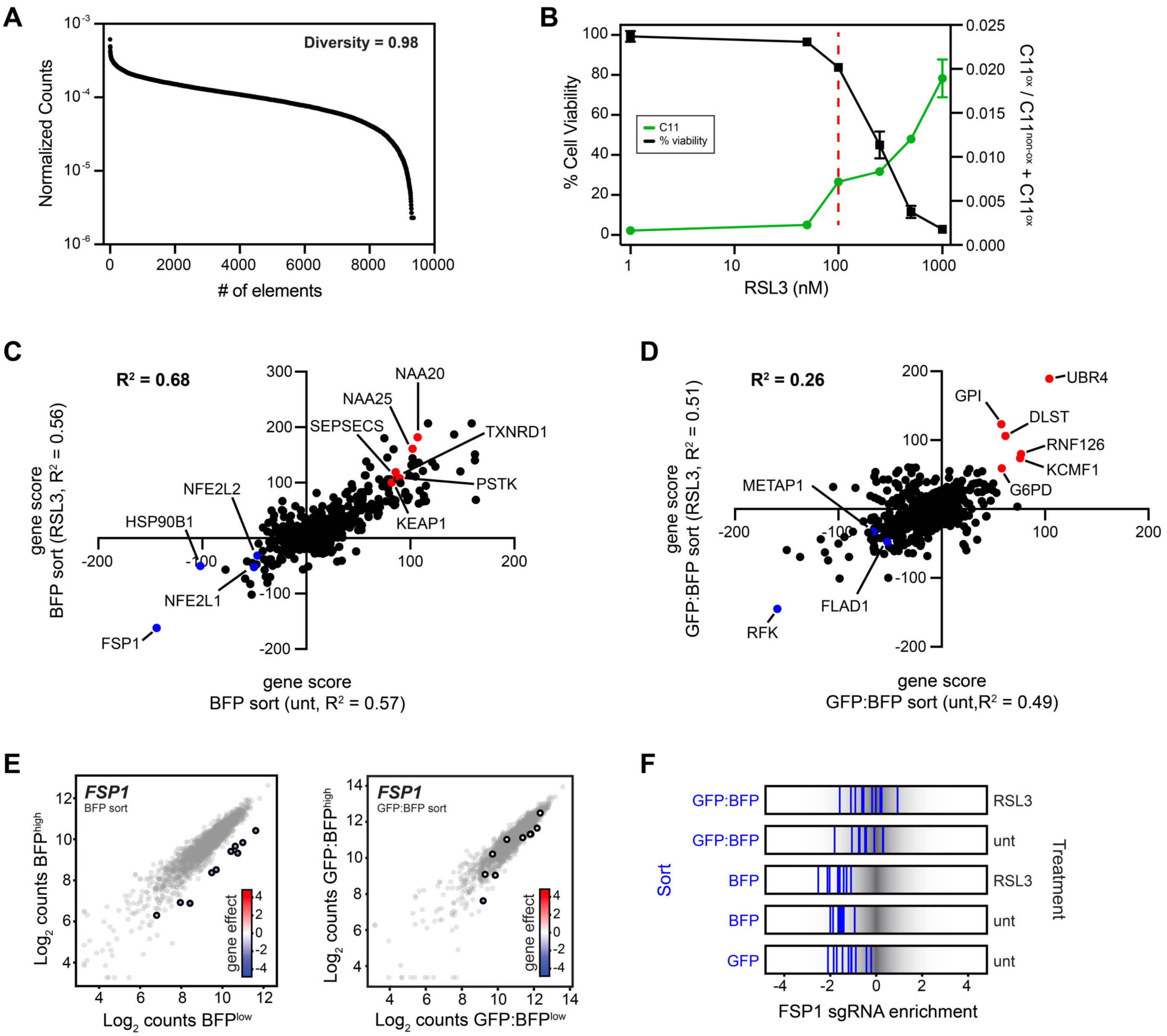

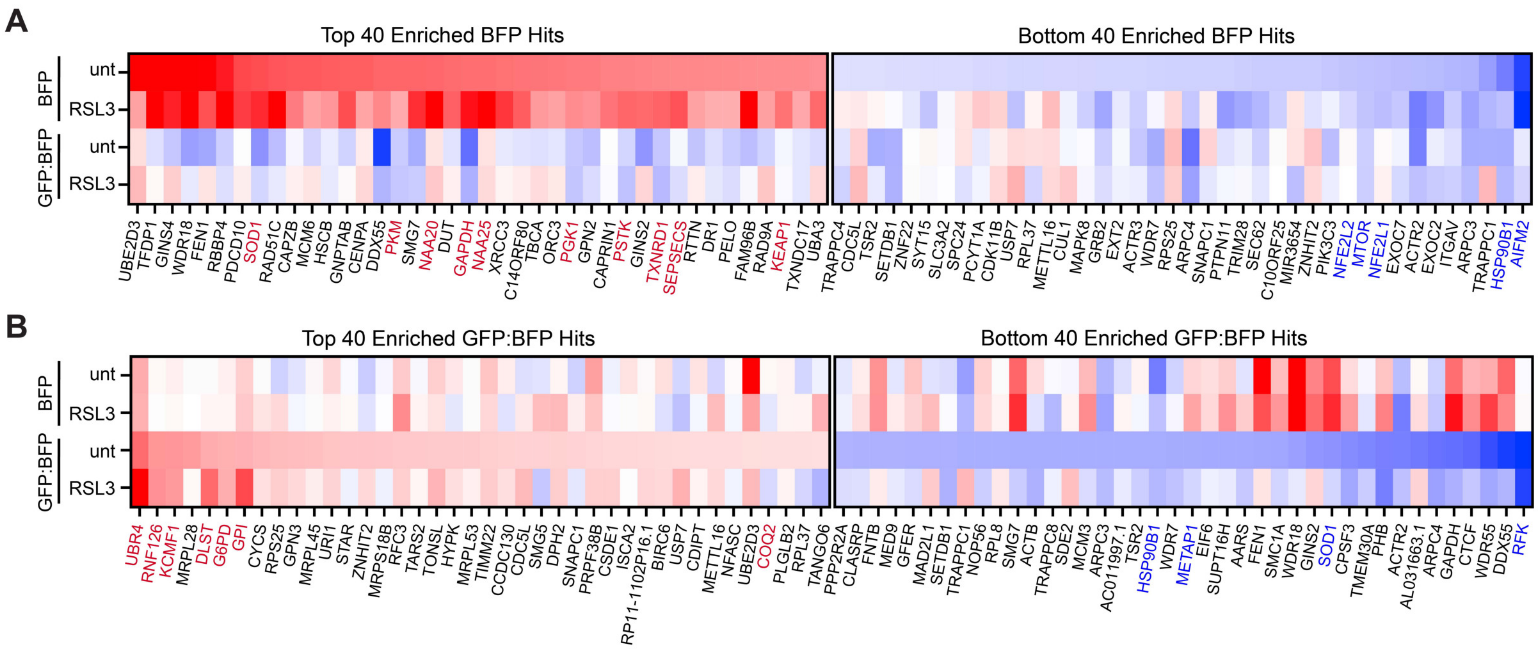

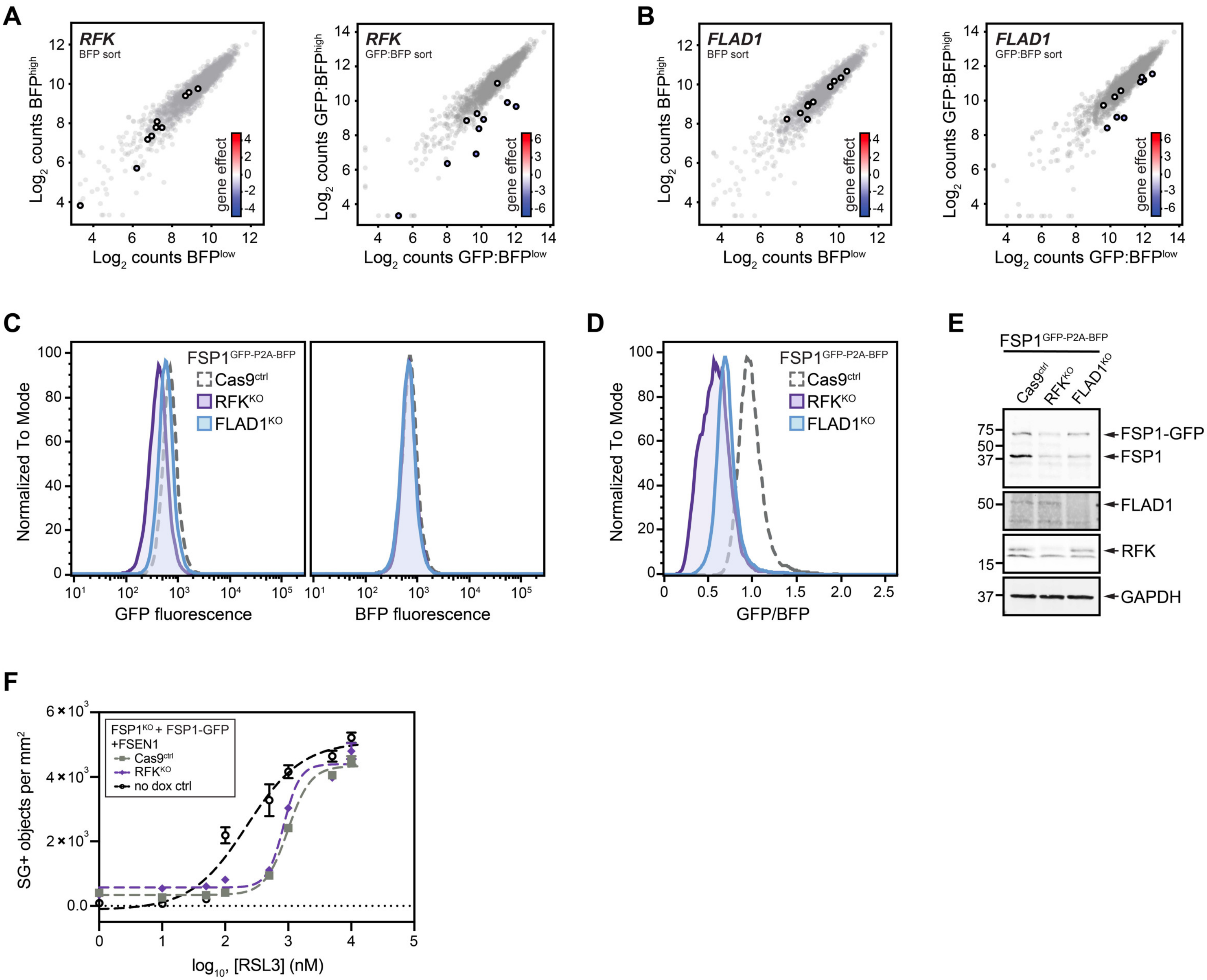

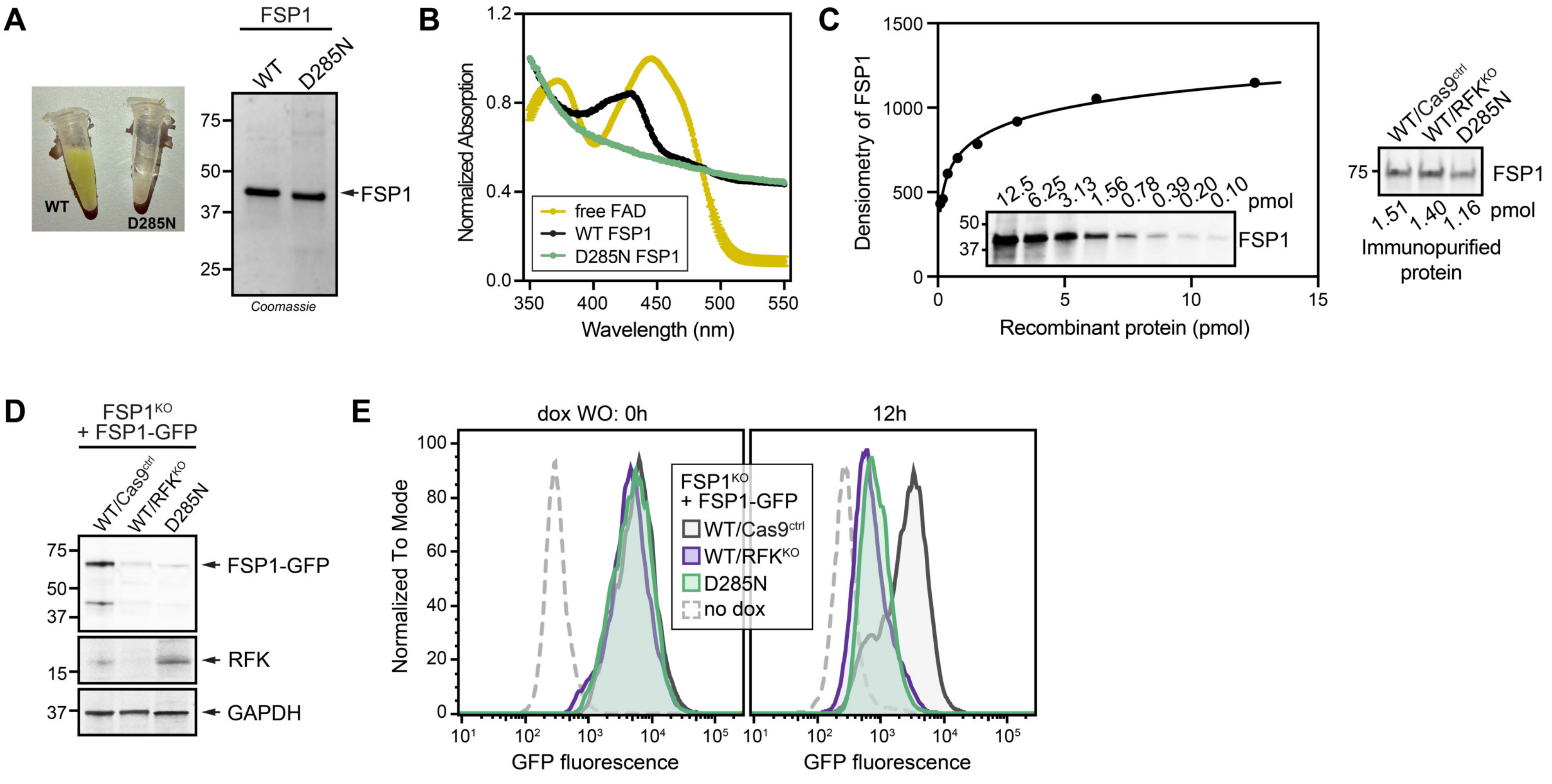

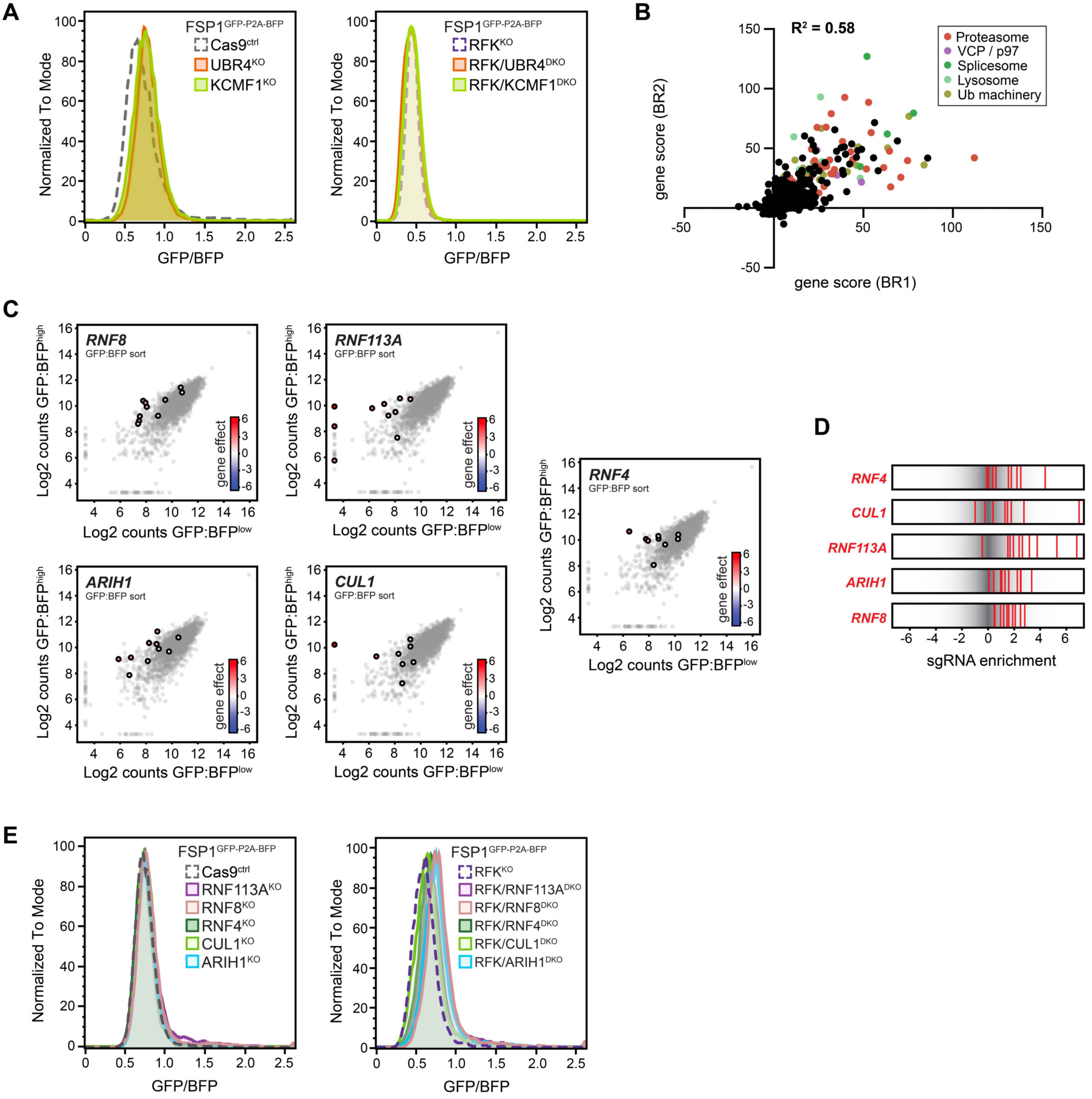

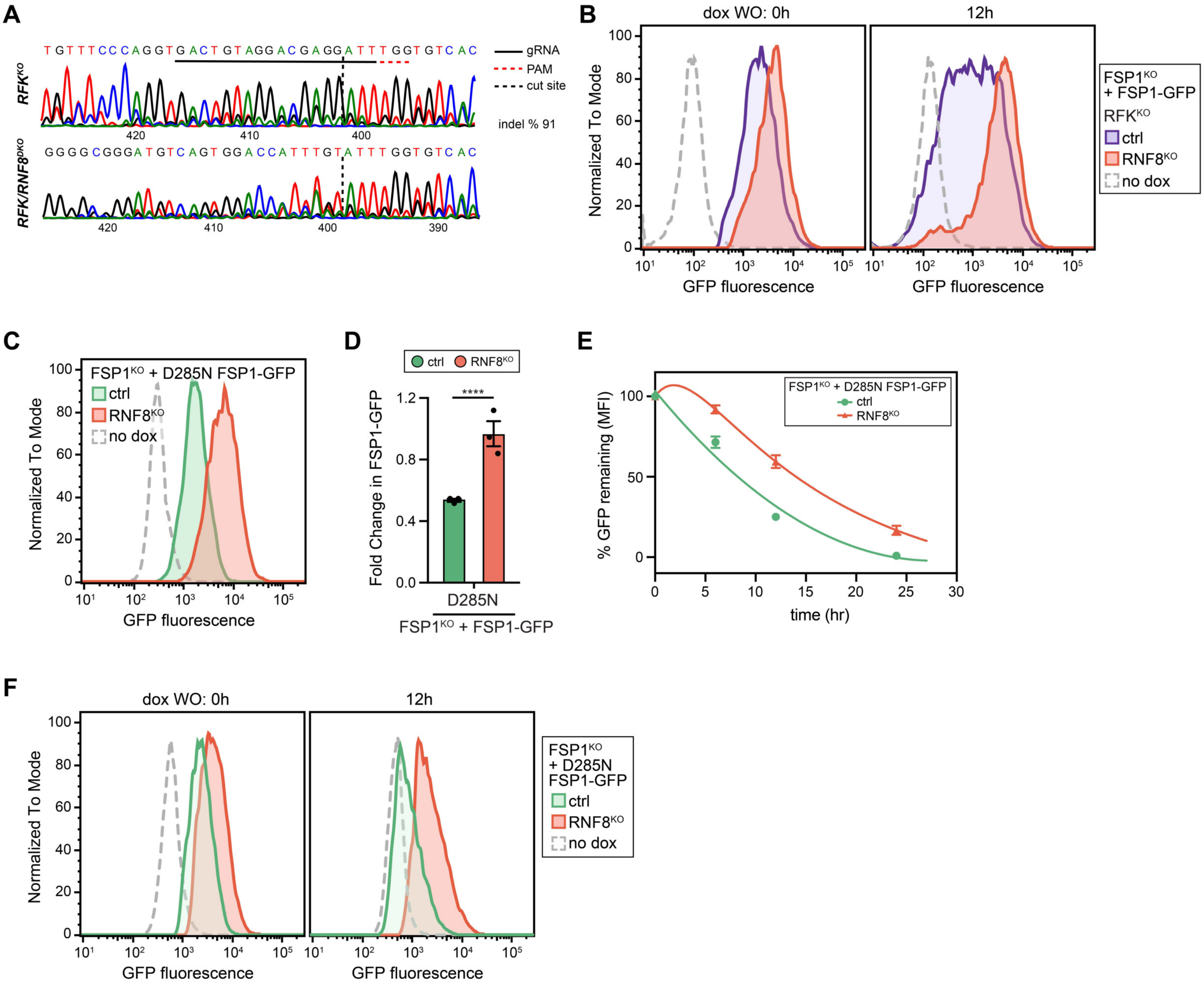
